# A quantitative approach to discover nonlinear signaling linking combinatorial environments with cellular responses

**DOI:** 10.1101/2025.10.21.683831

**Authors:** Dmitry Kuchenov, Frederik Ziebell, Florian Salopiata, Alix Thomas, Mevlut Citir, Ursula Klingmueller, Wolfgang Huber, Carsten Schultz

## Abstract

Cells constantly integrate diverse inputs from the extracellular environment, yet our understanding of how cells effectively process information from multiple cues at the same time remains limited. We utilized an integrated imaging platform and RNAseq analysis to investigate the combined effects of growth factors on cellular signaling and gene expression. Paired stimuli by receptor ligands revealed diverse signaling signatures— ranging from antagonism to synergy—driving global signaling programs and gene expression. Notably, correlation networks based on signaling signatures identified vulnerabilities in cancer cells when compared to synergistic drug combinations. Profiling kinase and phosphatase activities uncovered a crucial interplay, where cellular sensitivity and phospho-turnover dynamics are modulated by input history through coordinated basal kinase and phosphatase activities. Our novel methodology sheds light on cellular processing of multiple cues, elucidating intricate mechanisms underlying cellular adaptation to extracellular environmental variations.

Based on previous preprint: **doi:** https://doi.org/10.1101/346957

## Introduction

The extracellular microenvironment is a signal-rich system defined by cell-cell contacts and the presence of signaling molecules like nutrients, growth factors (GFs), cytokines, and hormones^1,2^. GFs and their receptors (receptor tyrosine kinases, RTKs) are well known to trigger a network of signalling pathways to govern diverse fate decisions and biological processes such as proliferation, apoptosis, differentiation and metabolism. Aberrant RTK signaling and downstream network malfunction is frequently involved in diseases including cancer^3–5^. Upon activation, RTKs at the plasma membrane induce signaling through a set of well-known intertwined signalling pathways that are shared among different RTK signalling networks^5–9^. Remarkably, despite a limited number of intracellular signaling components involved in RTK signaling, different RTKs have quite distinct functions *in vivo*^10,11^. While numerous studies have revealed fascinating signaling mechanisms of individual RTKs^6–8,12,13^, there is still a significant gap in understanding of: 1) how cells integrate and process information from multiple RTKs; 2) how they integrate this input with the history of microenvironment changes, and 3) how combinatorial RTK signaling is translated to gene expression patterns and in turn to physiological responses. Remarkably, combinations of GFs/cytokines may induce a unique gene expression profile as well as a distinct physiological outcome^14–18^. Such combinatorial stimulation schemes serve as a simplified models for the complex scenario provided by the extracellular microenvironment.

Multiple reports proposed that signaling signatures such as additivity, synergy and antagonism make important contribution in combinatorial signal processing^19–21^. Those signaling interactions stem from signaling cross-talk or feedback loops between and within pathways of the RTK network. Our work has also suggested that the concentration of GFs in a pairwise stimulation will program signaling signatures^22^. Such signatures might be unique and are not predictable from the individual components of a signaling network and/or the individual treatments with a single cue which makes signaling network analysis a challenge^20,23,24^. We suggest that a combinatorial stimulation with simultaneous quantitative monitoring of multiple signaling network events over time and gene expression profiling would help to better understand the integration of information from multiple RTKs and subsequently the function of a signaling network. Such extended data sets will reflect the complexity of the cellular signaling networks governing gene expression and cellular responses, but are labor-intensive to generate in a comprehensive fashion by conventional approaches. To meet this challenge, we utilize a FRET-based multi-parameter imaging platform (FMIP) that allowed monitoring the activity of 40 signalling molecules at the single cell level with high temporal resolution in a single experiment (**Figure 1A**)^22^.

**Figure 1.**
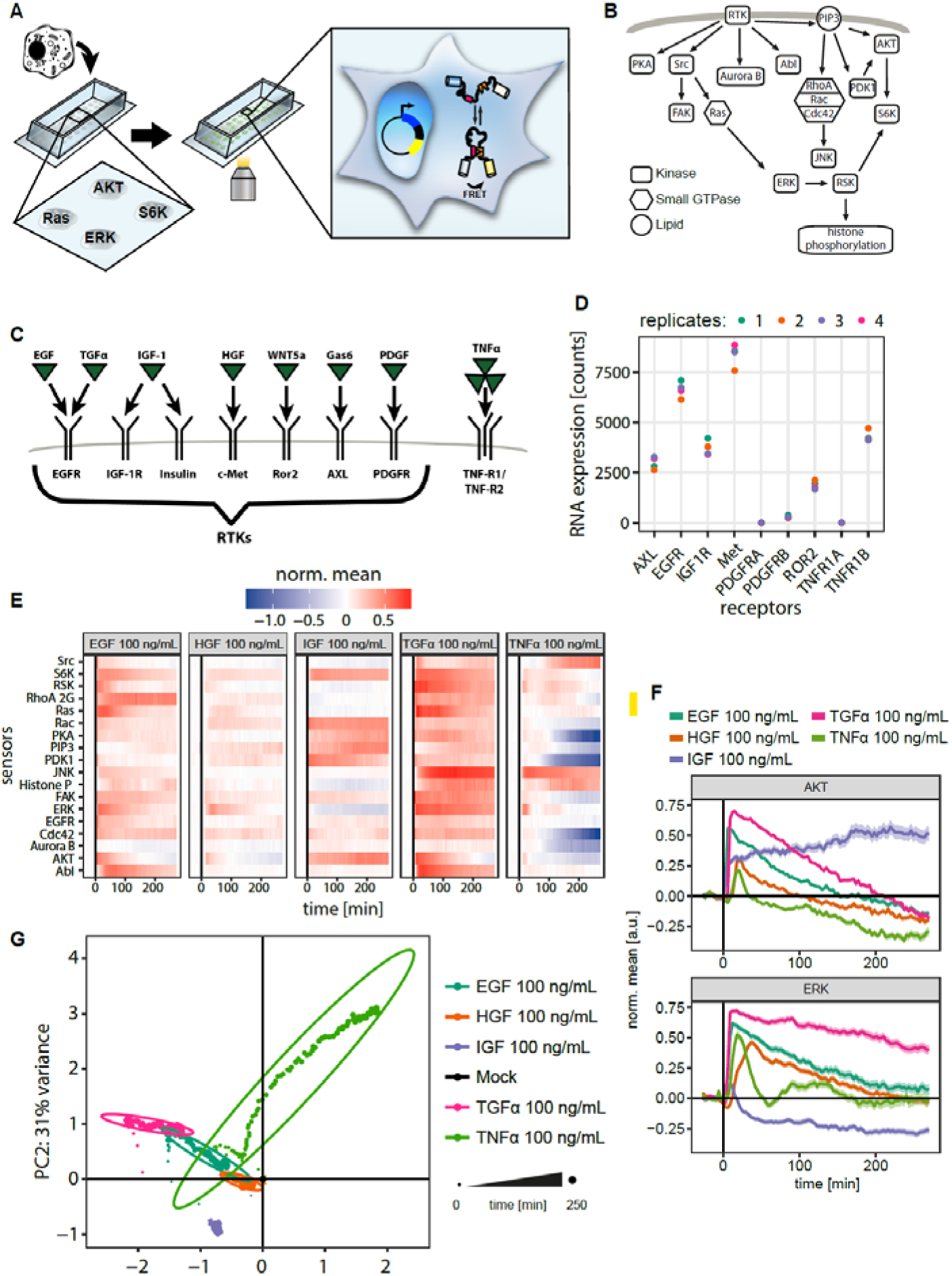
Dynamic signaling program is defined by cue identity. (A) Workflow of the FMIP assay. Plasmids encoding FRET biosensors were individually reverse transfected into human cells. FRET ratio was measured with an automated wide-field microscope. (B) Overview of the GF responsive signaling network measured by the FMIP. (C) Growth factor/cytokine receptors investigated in this work. (D) mRNA expression levels of the investigated receptors determined by RNA-seq after 16 h of serum starvation in HeLa cells. n = 4 independent experiments. (E) Dynamic signaling response after exposure to five different ligands. Color scale indicates mean FRET ratio over 270 min, normalized to the absolute maximum value (relative activity). (F) Representative examples of distinct dynamics over time after growth factor or cytokine treatment for the data quantified in (E). Curves indicate mean ± SEM. (G) Principal component analysis (PCA) of the average response to growth factors and cytokines. Each dot represents a single time point while the ovals indicate 95% of time points of the same treatment.

In this work, we show that pairwise GF stimulation (EGF, IGF-1, HGF, TGFα) shapes the global signaling program through dynamic signaling signatures (additivity, synergy, and antagonism). Notably, we observed that signature-based correlation networks provide a framework to identify synergistic drug combinations. Integrative analysis revealed that each signaling program correlates with unique gene expression profiles. We further demonstrate that in the presence of serum or combination of GFs, i.e. under quasi-physiological conditions, the cellular signaling network is pre-activated and tuned to achieve a unique signaling program in response to physiologically relevant concentrations of EGF. We also report that the basal activity of kinase and phosphatase that tunes phospho-turnover rate and phosphorylation status of downstream RTK effectors differs qualitatively and quantitatively depending on the presence of multiple signaling cues. We observe that the phospho-turnover rate adjusts signaling responsiveness of the cell and shapes physiological outcome for new incoming signal. Our methodology supports the hypothesis that a complex environment, defined by the identity and concentration dependent combination of extracellular cues, plays a crucial role in history-guided cellular responses to the altered environment of cancer cells.

## Results

### Quantitively analysis of TGFα, EGF, IGF-1, and HGF signalling differences

We employed a FRET biosensor-based multi-parameter imaging platform (FMIP) for monitoring 40 signalling events at the single cell level with high temporal resolution in a single experiment (**Figure 1A**)^22^. FMIP relies on transiently expressed FRET-biosensors to visualize enzyme activities in live cells rather than quantifying protein posttranslational modifications^25,26^. This platform quantitatively assesses the functional status of a signalling network.

To explore the mechanisms underlying the integration of multiple extracellular signals, our initial objective was to evaluate the signaling response induced by various growth factors. We stimulated HeLa cells with 100 ng/mL of platelet-derived growth factor BB (PDGF-BB), epidermal growth factor (EGF), tumor growth factor alpha (TGFα), insulin-like growth factor (IGF-1), hepatocyte growth factor (HGF), growth arrest-specific protein 6 (GAS6), wingless and int-related protein 5a (WNT5a), or tumor necrosis factor alpha (TNFα) and monitored response of 40 FRET biosensors by the FMIP (**Figure 1A**). We included TNFα, which binds to receptors within the TNFR protein family, to elucidate fundamental principles of signal integration for different receptor family. Each of these ligands binds to a well-known receptor inducing signaling that has been intensively characterized before (**Figure 1B-C**)^9,27^. Of the 40 FRET sensors, we identified 18 that reported consistent FRET ratio changes across independent replicates (n ≥ 2, ∼262 cells in average) under at least one condition and CV (Coefficient of Variation)<30% across all conditions tested (**Figure S2A**). Collectively, these responsive biosensors visualized diverse pathways of the signalling network including Ras/ERK, PI3K/AKT, JNK, and Src/FAK pathways (**Figure 1B**). In these cells, PDGF, Gas6 and Wnt5a were inactive or induced weak response and were excluded from further experiments. We detected immediate and strong signaling responses upon stimulation of cells with EGF, TGFα, IGF-1, HGF and TNFα (**Figure S1A**). Additionally, we confirmed the expression of corresponding receptors (EGFR, IGFR-1, Met and TNFRs), by measuring the abundance of mRNA in FBS starved HeLa cells (**Figure 1D**). Our quantitative analysis of signaling activity identified differences in the dynamics of signaling network activities between all five growth factors (**Figure 1E**). For example, in agreement with previous studies, TGFα induced much stronger signaling responses in comparison to EGF although both bind to the same receptor, EGFR^28–30^ (**Figure 1F**). Although TGFα and EGF induced the same downstream effectors, stimulation with EGF or TGFα produced a distinct transient dynamic trend of AKT and ERK kinase activity (**Figure 1F**). In turn, IGF-1 induced strongly sustained AKT activity, but led to a weak ERK activity followed by a slight inhibition after 10 min which is in line with previous studies using translocation probes in HeLa cells^31^. In contrast, stimulation with HGF produced only relatively moderate transient activities of AKT and ERK while TNFα induced strong bi-phasic ERK activity and weak transient AKT activity. We used principal component analysis (PCA) to visualize the time-resolved differential activation of signaling pathways (**Figure 1G and S1B**). The PCA plot shows that time points belonging to the same treatment form well-separated clusters, indicating strong differences in global signaling dynamics. This separation was even more apparent in a t-SNE visualization when compared to PCA (**Figure S1C**). Importantly, these visualizations (PCA and tSNE) show that the effects of EGF treatment were highly similar to those of TGFα in comparison to other treatments, suggesting a similarity of the signaling program induced by growth factors binding to the same receptor (**Figure 1G and S1C**). We confirmed for the cells used here that extracellular cues such as GFs and cytokines are able to induce distinct dynamic signaling programs.

### Framework analysis to quantify pairwise growth factor interactions that are non-additive

Following the hypothesis that two (or more) signaling cues can result in a different signaling program than a simple additive effect of individual growth factors^14,15,17^, we reasoned that the signaling interaction between extracellular cues is not only dependent on the identity of the GFs but also on their concentration as was shown for EGF/IGF-1 interactions^22^. We therefore treated HeLa cells with three pairs of growth factors (TGFα/IGF-1, EGF/HGF and IGF-1/HGF) at various concentrations. The data on signalling network activity were combined with our previous EGF/IGF dataset obtained under the identical experimental setup in HeLa cells^22^. To assess cell-to-cell variation in the signalling response, we calculated the average coefficient of variation (CV) across all time points for each treatment/FRET biosensor pair. We found that 87% of FRET biosensor-treatment pairs had a CV <15% indicating strong consistency of single cell responses across different conditions (**Figure S2A**). Thus, the data confirmed the precision of our signaling network activity analysis. Principal component analysis showed a clear separation of the global signalling responses upon treatment with a different concentration of each pair of GFs (**Figure 2A-C**). Importantly, PCA illustrated that the shift of the cluster of time points belonging to the same treatment is not proportional to the concentration or to the ratio of GFs for all GF pairs tested. This result is in line with previous findings^12,22^. Notably, although EGF and TGFα bind to EGFR^30^ they induced distinct signaling programs (**Figure S2B**) and also exhibited different profiles of signaling interaction with IGF-1, suggesting that the interaction is also tuned by the identity of the cue due to its unique signaling characteristics (**Figure 1E, 1F** and **S2B**). Because TNFα binds to the receptor of a TNFR protein family (distinct from RTK family), we measured signaling interaction between EGF/TNFα and IGF-1/TNFα to confirm general principles of signal integration for different receptor family. The analysis of signaling interaction between the pairs EGF/TNFα and IGF-1/TNFα revealed a clear shift in PCA space in response to a combinatorial treatment, suggesting similar mechanisms of signal integration based on synergy/additivity/antagonism profile (**Figure S2C**). As an independent confirmation that non-additive signature of growth factors is a common signaling feature for distinct cell lines, we observed that signaling signatures between EGF and IGF-1 also occurred qualitatively in human H838 non-small cell lung carcinoma cells (**Figure S2D**). Overall, these findings suggest that non-linear signal integration is a property of different cell lines and observed across various receptor families.

**Figure 2.**
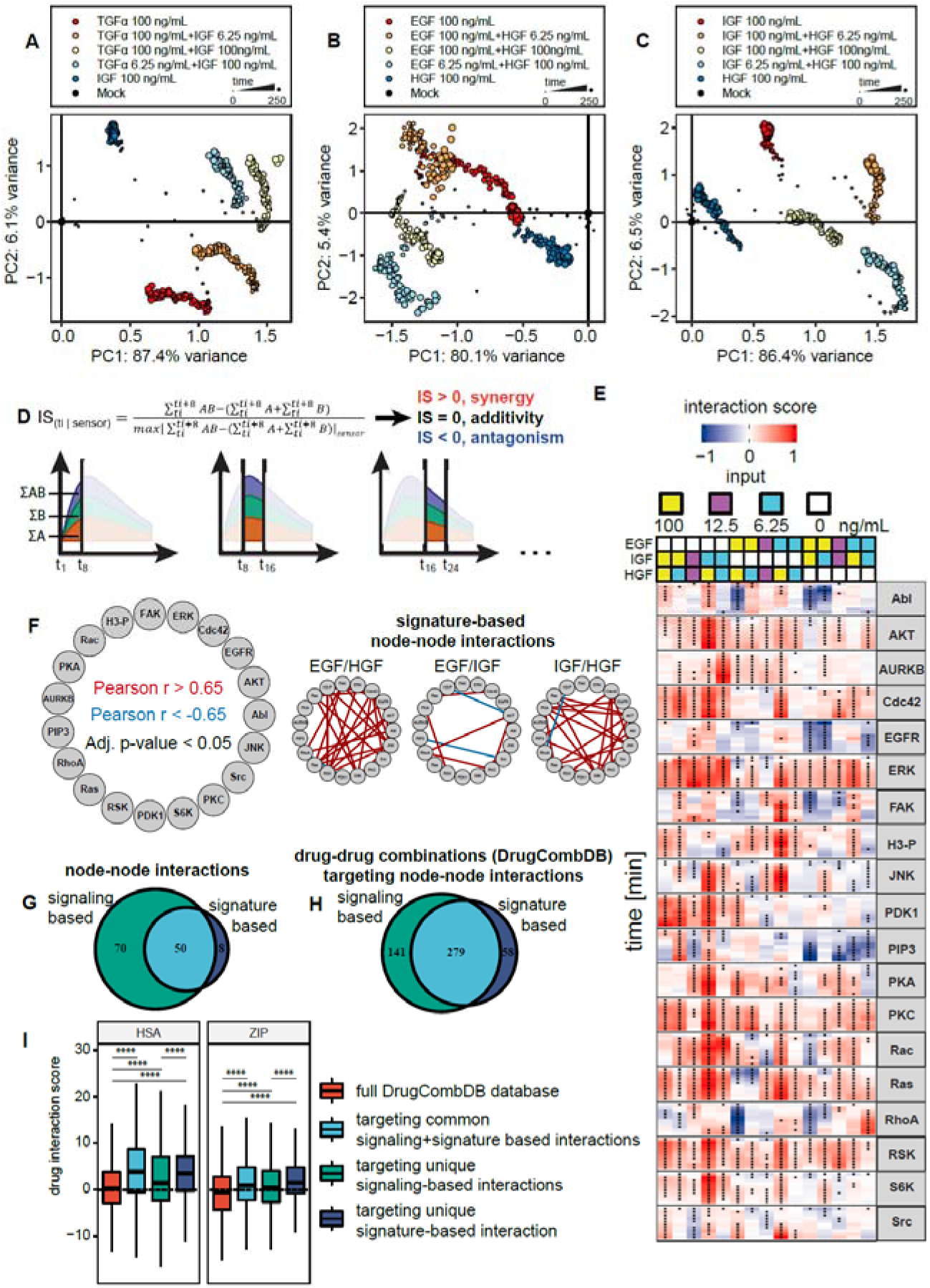
Signaling signatures induced by growth factor combinations identifies cancer vulnerabilities. (**A-C**) Global signaling response to the combinations of three ligan pairs projected into the first two principal components: (A) TGFα/IGF-1, (B) EGF/HGF, (C) HGF/IGF-1. Each dot represents a time point. Size indicates the time after treatment in min. (D) Calculation of the interaction score (IS) between growth factors. (E) The dynamic interaction ma reflects antagonism (blue), additivity, and synergy (red) over time in response to stimulation with a pair of GFs. Signaling interaction is quantified and classified as depicted in (D). All significant IS values (adjusted p-value < 0.05, Student’s two sample t-test) are marked with a black dot. (F) Graphic representations of strong Pearson correlations between signature profiles for EGF/IGF-1, EGF/HGF and HGF/IGF-1. Red and blue edges represent positive (r > 0.65 with adjusted p-value < 0.05, Student’s t test) and negative (r < -0.65 with adjusted p-value < 0.05, Student’s t test) correlations, respectively. (G) Venn diagram of node-node interactions identified across signature- and signaling-based correlation network. (H) Venn diagram of drug combinations targeting node-node interactions identified across signature- and signaling-based correlation network. (I) Boxplot of synergy scores (HAS and ZIP) found in the entire DrugCombDB database, drug combinations targeting common signaling- and signature-based interactions, drug combinations targeting unique signaling-based interactions and drug combinations targeting unique signature-based interactions

To gain deeper insights into the mechanisms governing distinct signaling programs when two growth factors are present and to quantify non-additive signaling responses, we introduced an interaction score. An interaction score (IS, a Benjamini-Hochberg adjusted p-value < 0.05 see Materials and Methods) quantifies the difference, within a specified time window, between the response to the combined stimuli A and B and the sum of responses to the individual stimuli. This score is then normalized by dividing it by the maximum absolute IS value observed for each FRET sensor (**Figure 2D)**. The IS detects non-additive signaling interactions (synergy and antagonism) but remains largely unbiased with regard to the underlying molecular mechanisms. A positive IS value indicates synergistic behavior between stimulus A and B whereas a negative IS value suggests antagonistic or saturating behavior. At the same time, IS = 0 represents an additive response or the absence of a response. First, we confirmed that the most common interaction type was additive (IS distribution is centered at zero) (**Figure S2E)**. This observation is also in line with a finding suggesting that additive interactions are dominant within signaling network^21^ indicating that our multiplicative normalization worked well over the full range of observed synergy and antagonism. We then created a composite interaction map to visualize signaling interaction over time between each pair of growth factors (**Figure 2E** and **S2F**). Notably, the interaction mode between two extracellular cues was highly dynamic which is consistent with the PCA of the global signaling response (**Figure 2A-C** and **S2B-C**). All growth factor pairs displayed a distinct profile of IS, supporting previous observations^20,22,23^. Importantly, for each pair of GFs, the signaling interaction mode was dependent on the concentration of the two stimuli in a combination explaining the non-linear shift in the PCA space for each growth factor pair.

A major motivation for quantitatively measuring and systematically mapping of the IS between a pair of GFs is the possibility of identifying unanticipated cross-talk between growth factors. We hypothesized that signaling molecules with highly correlated IS profile tend to be directly or indirectly co-regulated under pairwise treatment. Such signaling events might be missed by looking at raw signaling dynamics. To assess GF pair-specific signaling interactions, we computed pairwise Pearson correlations of time-dependent IS patterns for each GF pair (EGF/IGF-1, EGF/HGF and HGF/IGF-1), a signature-based approach, as well as of raw signaling time courses for each condition (EGF, IGF, HGF, EGF/IGF-1, EGF/HGF and HGF/IGF-1), a signaling-based approach, for all measured FRET biosensors. Next, we computed a correlation network to visualize the architecture of signature-based and signaling-based interactions for each growth factor pair or condition (**Figure 2F and S2G**), where we show the pair of sensors with significant correlations (|r|>0.65, a Benjamini-Hochberg adjusted p-value < 0.05 by Student’s two sample t-test). Although we observed strong overlap, the analysis revealed that the signaling-based approach identified more node-node interactions (120 interactions) in comparison to a signature-based network (58 node-node interactions) (**Figure 2G)**. For example, we detected positive IS profile correlation between JNK and S6K (**Figure 2F and S2G**) by signalling- and signature-based correlation approaches. In support of this notion, it was previously shown that active JNK1 is able to directly phosphorylate S6K under various experimental settings^32–34^. Notably, only signature-based analysis identified positive correlation between Src and PKC. This is in line with the facts that Src physically interacts with atypical PKC in vivo^35^ and Src phosphorylates PKC delta^36^ suggesting that at least some of the detected signature-based correlations may be a result of direct interplay between two signaling molecules that is not detected by the signaling activity correlation approach.

We next asked whether the identified node-node interactions are physiologically relevant for cell proliferation and survival. Massive drug combination data in the context of cancer cell proliferation and survival can now be used to study the outcome of targeting node-node interactions. Using DrugcombDB data^37^, we could identify 478 unique drug-drug combinations targeting signaling- and signature-based node-node interactions (**Figure 2H**). Drug synergy analysis suggests that drug combinations targeting node-node interactions identified by both signaling- and signature-based approaches show stronger synergistic effect in comparison to total DrugcombDB database (HSA: p-value < 2.22x10^-16^ and ZIP: p-value = 1x10^-12^ by Student’s two-sided t-test) (**Figure 2I**). Importantly, we observed higher HSA and ZIP scores (HSA: p-value = 1,3x10^-10^ and ZIP: p-value < 2,22x10^-16^ by Student’s two-sided t-test) for drug combinations targeting unique node-node interactions identified by signature-based approach only in comparison to drug combinations targeting unique signaling-based node-node interactions. Thi suggests that the majority of observed signature-based interactions are physiologically relevant in the context of different types of cancer cells and involved in regulation of proliferation and survival.

### Integrative framework to identify gene expression program driven by combinatorial treatment

Assessing the impact of a signaling program on a gene expression profile is fundamental in understanding the link between extracellular environment and the physiological state of the cell. We therefore investigated whether concentration-dependent signaling programs of growth factor pairs were associated with changes in gene expression. We specifically focused on the EGF and IGF-1 combination since this pair induced distinct signaling programs (**Figure 1E-G**) and showed strong concentration-dependent signaling interactions (**Figure S2B** and **2E**). Using genome-wide RNA-seq, we quantified the mRNA abundance upon individual or combinatorial treatments in triplicates at 4 hours after stimulation (**Figure 3A**). We first confirmed that our RNA-seq analysis reproduced previously published data from HeLa cells treated with 20 ng/mL EGF^38^ (**Figure S3A**). Biological replicates for different samples gave highly similar transcription profiles (**Figure S3B**), confirming the accuracy and precision of our RNA-seq analysis. We further globally examined the effect of individual and combinatorial treatments on the gene expression profile. PCA showed a pattern of distinct gene expression profiles that were similar to the pattern of the signaling network states (**Figure S3C**) suggesting that concentration-dependent signaling programs result in a distinct transcriptional output. Importantly, co-inertia analysis^39^ confirms that gene expression states induced by differential combinations of EGF and IGF-1 follow signaling programs induced by the same EGF/IGF1 combinations (short arrows; RV coefficient 0.83), indicating that observed signaling and RNA-seq data have strong similarity in their global response and that the observed set of signaling molecules plays a significant role in the regulation of the global gene expression state upon GF treatment (**Figure 3B**).

**Figure 3.**
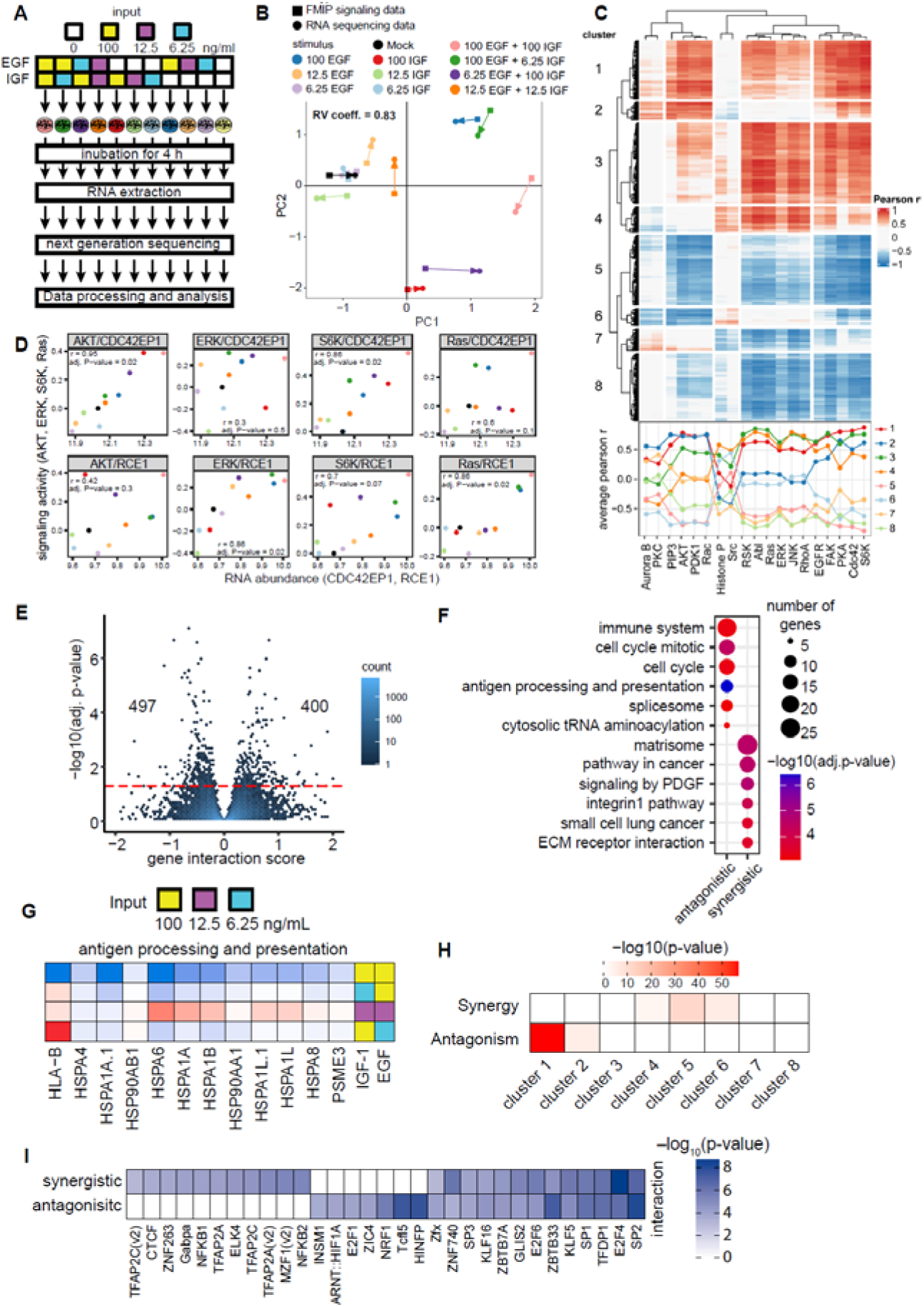
Combinatorial cytokine code governs gene expression profile through the signaling network activity state. (A) Experimental setup applied to determine combinatorial GF code-specific transcriptomes. HeLa cells were stimulated with various concentration combinations of EGF and IGF (individually or pairwise) for 4 h. Subsequently, cells were lysed, mRNA was extracted and subjected to RNA-seq analysis. (B) Co-inertia analysis of signaling data and RNA-seq data across different growth factor combinations and concentrations (arrow base: FMIP data; arrow head: RNA-seq data) showing that the information carried by signaling and RNA-seq data was closer to each other in the same condition than across conditions. Vectors represent the proportional divergence between data sets within each condition. Shape of the point are used to aid identifying data type. Color indicates condition. (C) Clusters of significant Pearson correlations (adj. p-value < 0.05, with at least one FRET sensor) across the activity of signaling events. Pearson correlation values were determined for the rlog-transformed counts (DEseq2) of each gene with respect to the mean of signaling dynamics for each sensor (area under the curve). Clustering was performed on the correlation values using hierarchical clustering based on DIANA (Divisive analysis) algorithm for gene clusters and Ward’s method with Euclidean distance for signaling clusters. (D) Representative gene expression correlations from clusters 2 and 4, illustrated by RCE1 and CDC42EP1, respectively. Plots show how gene expression levels vary in relation to the dynamic signaling activity—quantified as area under the curve (AUC)—for key signaling molecules: Ras, ERK, S6K, and AKT. These molecules represent distinct signaling module clusters highlighted in (C): (1) **ERK**/**RAS**/RSK/ABL/JNK/RhoA, (2) EGFR/FAK/PKA/Cdc42/**S6K**, and (3) PIP3/**AKT**/PDK1/Rac1. (E) Volcano plots showing the gene interaction score (GIS) for antagonistic and synergistic genes. Red dotted line indicate threshold with significant FDR (= 0.05). (F) Gene set enrichment analysis of antagonistic and synergistic genes that responded to at least one pairwise treatment (adjusted p-value < 0.05). (G) Heatmap visualizing gene interaction scores of genes assigned with antigen processing and presentation from (F). (H) Heatmap shows gene clusters from (C) that were significantly enriched in each interaction response gene (IRG) group (Fisher’s exact test). (I) Heatmap shows known transcription factor (TF) binding motifs that were significantly enriched in each interaction response gene (IRG) group (with adjusted p value < 0.05 and abs (gene interaction score (GIS)) > 0.5). TFs with p value < 0.001 and RNA expression > 500 counts were defined as significantly enriched.

The differential gene expression analysis identified 7369 genes that change their expression at least in one of the condition (DESeq2 method^40^, FDR= 0.05 and abs(log2FC) > 0.3) in comparison of treated against control cells. As expected from differential pathway activation (**Figure 1E** and **F**), the differentially expressed genes (DEGs) induced by individual treatments varied considerably between EGF and IGF-1, especially at low GF doses (**Figure S3D**). Next, we directly compared DEGs between each condition. We were able to identify that each condition also regulated large unique gene sets, except for individual treatments of the lowest (6.25 ng/mL) concentration (**Figure S3E and S3F**). Overall, these data suggested that concentration-dependent GF signaling programs result in distinct gene expression profiles.

We sought to further link the concentration-dependent signaling program with gene expression profile by integrating FMIP (FRET-based Multi-parameter Imaging Platform) and RNA-seq data. We calculated the correlations between the signaling response for each FRET sensor (area under the curve) and gene induction across the combinatorial growth factor treatments (**Figure 3C** and **3D**). Using an unsupervised clustering analysis, we identified 5 distinct signaling and 8 distinct gene clusters (**Figure 3C**). As expected, the PIP_3_/PDK1/AKT/Rac and Ras/ERK/RSK/JNK/RhoA/Abl signaling modules are clustered separately. Interestingly, we observed 2 symmetric gene cluster types that have opposite (positive and negative) correlation profile across signaling network (clusters 1/5, 2/6, 3/8 and 4/7) indicating that the same signaling program results in up-regulation and down-regulation of gene expression. Genes in cluster 3 and 4 were distinguished by strong correlations with the Ras/ERK/RSK/JNK/RhoA/Abl signaling module, with RCE1 as an example (**Figure 3C** and **3D**). As expected, we show that previously identified ERK-up-regulated genes^41^ are only enriched in cluster 3 and 4 (p-value = 1.95*10^-5^ and 8.5*10^-6^, respectively, Fisher’s exact test) indicating the robustness of our clustering approach. Importantly, an enrichment analysis for biological process shows that each gene cluster is associated with a unique set of biological functions (**Figure S3G**).

To gain insight into the unexpected molecular consequences that were triggered in response to the combinatorial EGF/IGF-1 treatment at the level of gene expression, we calculated a gene interaction score (GIS) to quantify non-linear transcriptional interactions induced by pairwise GF treatment. We identified 897 antagonistic and synergistic genes (**Figure 3E**, FDR < 0.1) that were then categorized in 9 groups according to the interaction profile similar to a recently published classification^15^ (**Figure S3H**). We found that the vast majority of interactions between EGF and IGF-1 were well approximated by an additive interaction, indicating that most gene expression changes are accurately predictable from individual treatments, which is in line with the observation that most genetic interactions are additive^42^ and similar to signaling interactions (**Figure S2E** and **S3H**). Interestingly, for synergistic and antagonistic GISs the predominant influence on gene expression was buffering, where individual treatments induced changes in the same direction and the paired stimulation also resulted in an effect in that direction, but lower than the sum (332 for synergistic buffering and 178 for antagonistic buffering, **Figure S3H-I**). Further analysis of biological process enrichment for interaction response genes (IRGs) that responded synergistically or antagonistically to at least one pairwise treatment (FDR < 0.05) shows overrepresentation of pathways linked to synergistic regulation of the extracellular matrix (ECM), focal adhesion, integrin, PDGF, and cancer related signaling (**Figure 3F**). We observed a significant enrichment for antagonistic IRGs involved in the regulation of mitosis, cell cycle, splicing and mRNA destabilization (**Figure 3F**). Overall, these data suggested that concentration-dependent GF signaling programs result in complex changes of the gene expression profile in additive and non-additive manner.

Whereas synergistic and antagonistic effects were observed under all tested concentrations of EGF and IGF-1, global analysis of IRGs suggested that the effects on individual genes differed profoundly for each condition (**Figure S3H**), mainly expressed as a different interaction profile that was concentration-dependent (**Figure S3K)**. For example, we observed strong concentration dependence of the interaction profile within the antigen processing and presentation gene ontology module (**Figure 3G)**. Interestingly, the synergistic effect in this module was observed under low concentrations of GFs (12.5ng/ml EGF + 12.5ng/ml IGF) whereas the high concentration of one of the GF in the combination resulted in an antagonistic effect. To further explore the link between gene interaction modes and signaling network activity, we calculated the enrichment of gene clusters (**Figure 3C**) within antagonistic and synergistic genes (**Figure 3E**). We observed strong enrichment of cluster 5 (p-value = 2.9*10^-12^, Fisher’s exact test) in synergistic genes and cluster 1 in antagonistic genes (p-value = 1.2*10^-56^, Fisher’s exact test) (**Figure 3H**). Those clusters correlate with 3 distinct signaling modules suggesting that synergistic and antagonistic gene interaction is controlled by a crosstalk between different signaling pathways.

To identify transcription factors (TFs) that decode the synergistic and antagonistic signaling and tune gene expression upon pairwise GF treatment, we searched for over-representation of TF binding motifs in promoter regions of the 119 synergistic and 88 antagonistic genes (|GIS| >0.5 and a Benjamini-Hochberg adjusted p-value < 0.05) using Pscan^43^ and the JASPAR non-redundant database^44^. We identified a significant enrichment (p-value <0.001) for sites binding to 31 TFs expressed in HeLa cells and classified them into three subgroups (**Figure 3I**): 11 TFs that exhibited the strongest enrichment in synergistic genes; 7 TFs that exhibited the strongest enrichment in antagonistic genes; and 13 shared TFs that were enriched in both. Importantly, among TFs enriched in antagonistic genes, we identified histone nuclear factor P (HiNF-P or MIZF), which interacts with methyl-CpG-binding protein-2 (MBD2) to promote DNA methylation and transcription repression^45^. Among TFs enriched in synergistic genes, we found TFAP2A and SP1, both known to be associated with gene transcription activation in HeLa cells^46^. Collectively, our data indicate that observed concentration-dependent signaling states result in a distinct gene expression profile tuned by non-linear transcriptional interactions and regulate a diverse set of cellular processes and functions.

### Physiologically relevant concentration shapes signaling and cellular response

The physiological concentration range of GFs are much lower than the concentrations that are typically applied in cellular signaling studies with serum-starved cells^6,29,47^. Our experiments with physiologically relevant concentrations (6.25 and 12.5 ng/mL) of growth factors revealed that pairwise treatments modulate the strength of signaling response depending on the identity and concentration of extracellular cues (**Figure 2A-C** and **2F)**. Furthermore, gene expression analysis demonstrated the boost of total DEGs upon co-treatment with EGF and IGF-1 of low concentration (12.5 ng/ml) (**Figure 4A** and **S4A**) indicating strong interaction of EGF and IGF-1 at low concentrations on the level of gene expression. Therefore, we reasoned that complex extracellular environment defined by the presence of multiple cues tunes the signaling network state. To study whether the extracellular environment leads to altered sensitivity and response programs, we employed 10% fetal bovine serum (FBS) or well-defined mixture of GFs to mimic complex extracellular environment and manipulate basal phosphorylation state of the cell. In thi scenario, we studied the dynamic signaling program upon stimulation with EGF at low and high concentrations in overnight serum starved and non-starved HeLa cells. As expected, the EGF-induced signaling program was altered in the presence of 10% FBS (**Figure 4B**). Notably, the signaling response after a low concentration of EGF (6.25 ng/ml) treatment was much stronger in the presence of FBS compared to serum starved HeLa cells (**Figure 4B-D**). For example, we observed a stronger response of Src, ERK, and S6K in non-starved cells (**Figure 4C**). Immunoblot analysis qualitatively confirmed that the presence of 10% FBS elevated EGF-induced pS6 levels, confirming the FRET data (**Figure 4D** and **S4B**). Next, we analyzed the global signaling response by PCA in serum starved and non-starved cells. We observed a clear separation between serum-starved cells treated with a range of EGF concentrations and non-starved cells, indicating that the presence of 10% FBS tunes the basal activity of the network to respond to the EGF stimulation (**Figure 4E**). Importantly, we observed strong differences induced by low concentration of EGF between starved and FBS-supplied cells that corresponds to the strong shift in the PCA space. In contrast, PCA showed that the presence of FBS resulted in a relatively modest effect on the signaling responses when cells were stimulated with a high concentration of EGF which is in line with a previous report^48^ (**Figure 4E**). By precisely controlled experiments where cells were pretreated with 12.5 ng/ml IGF + 12.5 ng/ml HGF, we also showed a distinct response to the low concentration of EGF (6.25 ng/ml) in comparison to starved cells, suggesting that the observed effect is relevant for physiological growth factor concentrations (**Figure S4C**). Together, these data indicate that the presence of multiple extracellular cues considerably tunes the signaling dynamic program in response to the GF stimulation of low concentration. This has major implications for comparing results of starved and non-starved cells in cell biology experiments.

**Figure 4.**
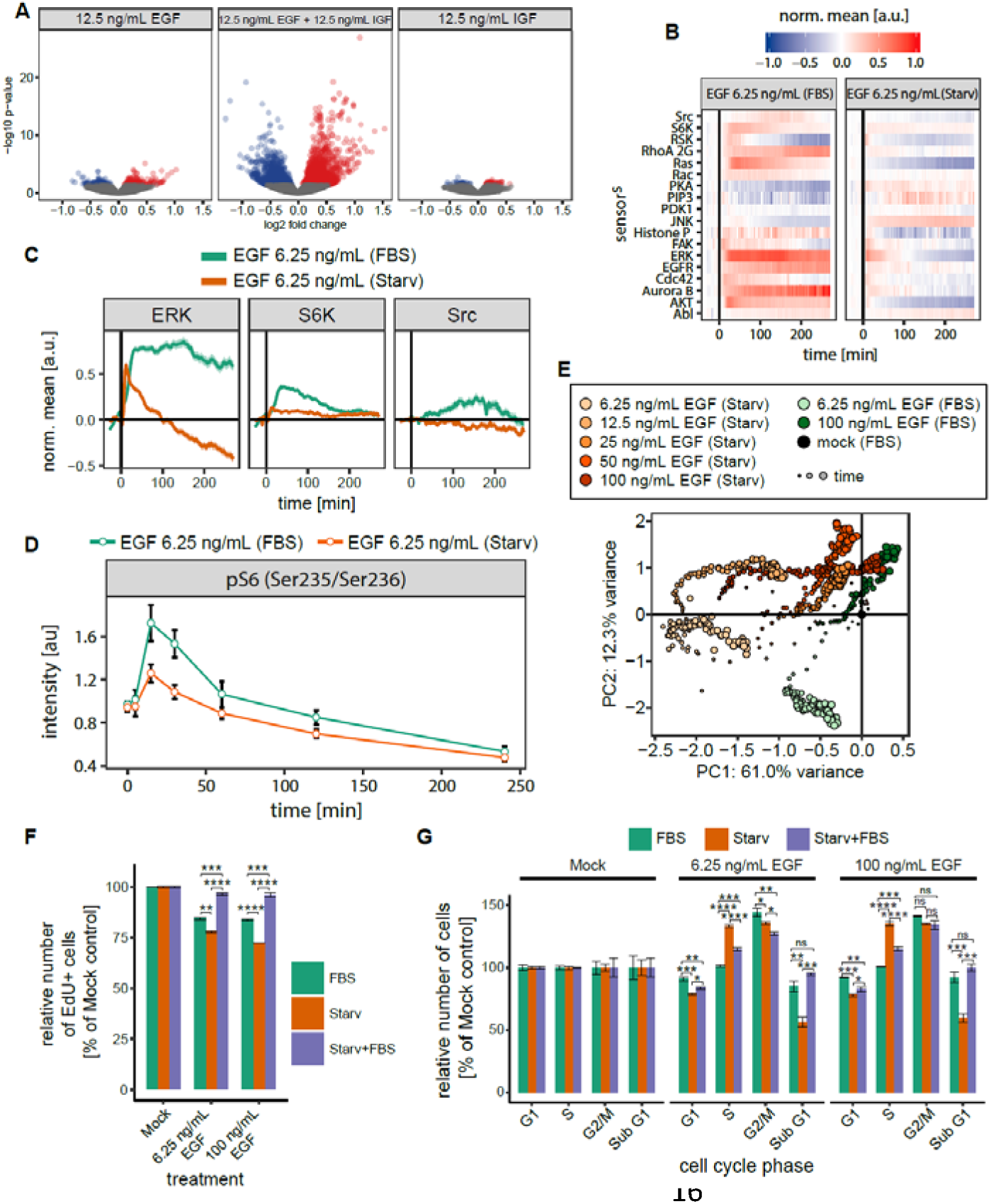
Quasi-physiological condition potentiates and shapes signaling response. (A) Volcano plots comparing cells treated with 12.5 ng/mL EGF, 12.5 ng/mL IGF-1 and 12.5 ng/mL EGF + 12.5 ng/mL IGF-1. Data represent three independent experiments. Blue and red dots indicate suppressed and induced genes with adjusted p value < 0.05, measured by DESeq2, respectively. (B) Quantification of the dynamic program of signaling activity after exposure to 6.25 ng/mL EGF in presence or absence of 10% FBS. (C) Examples of potentiation of signaling events in the presence of 10% FBS from (B). (D) Phosphorylation of S6 detected by quantitative immunoblotting. HeLa cells were maintained in either growth factor depleted or 10% FBS supplemented medium and stimulated with 6.25 ng/mL EGF. Error bars represent ± SEM. (E) Global signaling response to a range of EGF concentrations projected into the first two principle components. (F) Relative 5-bromodeoxyuridine (EdU) incorporation for cells kept in 10% FBS or serum starved cells. Cells were left unstimulated (mock) or stimulated with 6.25 ng\mL or 100 ng\mL EGF. Error bars represent ± SEM. (G) Comparison of the relative change of each cell cycle phase (G1, S and G2/M) between cells kept in 10% FBS supplemented or serum deprived medium. Cells were left unstimulated or stimulated with 6.25 ng/mL or 100 ng/mL EGF. Error bars represent ± SEM. Significance was assessed by Student’s t test. ns: p > 0.05, *: p ≤ 0.05, **: p ≤ 0.01, ***: p ≤ 0.001, ****: p ≤ 0.0001.

To determine the physiological relevance of signaling difference induced by the presence of 10% FBS, we quantified proliferation and determined cell cycle parameters assayed by 5-ethynyl-2-deoxyuridine (EdU) incorporation and 4,6-diamidino-2-phenylindole (DAPI) staining, respectively. As expected, serum starvation resulted in a reduced basal proliferation as well as increased fraction of cells in G1 phase (**Figure S4D**). Stimulation with EGF resulted in the inhibition of cell proliferation as well as an increase in the percentage of cells in G2/M phase regardless of the starvation status, a well-known effect of EGF on cancer cell lines^49–51^ (**Figure 4F-G** and **S4D**). This finding is also in line with the fact that mitotic cell cycle genes were highly enriched in the antagonistic gene set (**Figure 3F**). We further quantified that HeLa cells subjected to starvation and treated with EGF demonstrated a stronger decrease in proliferation in comparison to cells treated with EGF in the presence of 10% FBS (**Figure 4F**). Importantly, we observed that there is no difference in proliferation of starved cells between 6.25 ng/ml and 100 ng/ml of EGF suggesting that the effect of EGF is already saturated at the physiological concentration. EGF treatment of starved cells resulted in a significant increase in the relative number of cells in S phase compared to cells kept in full media. The latter showed a strong decrease in the number of cells in G_1_ phase (**Figure 4G**). Collectively, these data suggest that the molecular composition of FBS tunes the signaling network state: 1) to achieve higher signaling responsiveness of the cells and 2) to shape the EGF-induced signaling program and physiological response under physiologically relevant EGF concentrations.

### A quantitative framework to studying the basal signaling state of kinases and phosphatases

Although, it has long been known that serum is able to potentiate EGF- and IGF-1-induced responses^52,53^, the molecular mechanisms of signaling network tuning and pre-activation by multiple extracellular cues have not been explicitly identified yet. Recent work suggested that the processing of information from multiple cues might occur at the receptor level^12^. To address this possibility, we quantified total and plasma membrane EGFR expression by immunoblot analysis and immunostaining, respectively. Although we observed a slight increase in the expression of plasma membrane EGFR upon starvation (**Figure 5A** and **5B)**, we did not detect a significant difference in total EGFR as well as in ABL and SRC expression between cells kept in full media and serum starved cells (**Figure S5A, S5B and S5C**). Our findings indicate that tuning the signaling program by the extracellular environment occurs within the intracellular signaling network, i.e. by adaptation of the basal activity of downstream components. We therefore compared the basal activity state of the signaling network in the presence or absence of 10% FBS. Interestingly, basal signaling profiling revealed that all signaling events in our data set exhibited a high variability at the single cell level. As expected, we found that basal patterns were classified into three groups: signaling events either showed a decrease, an increase or no difference in basal activity upon starvation (**Figure 5C**) and compared well to a previous study^54^. This further supports our hypothesis that a signaling network is tuned by extracellular conditions to achieve a specific basal activity state.

**Figure 5.**
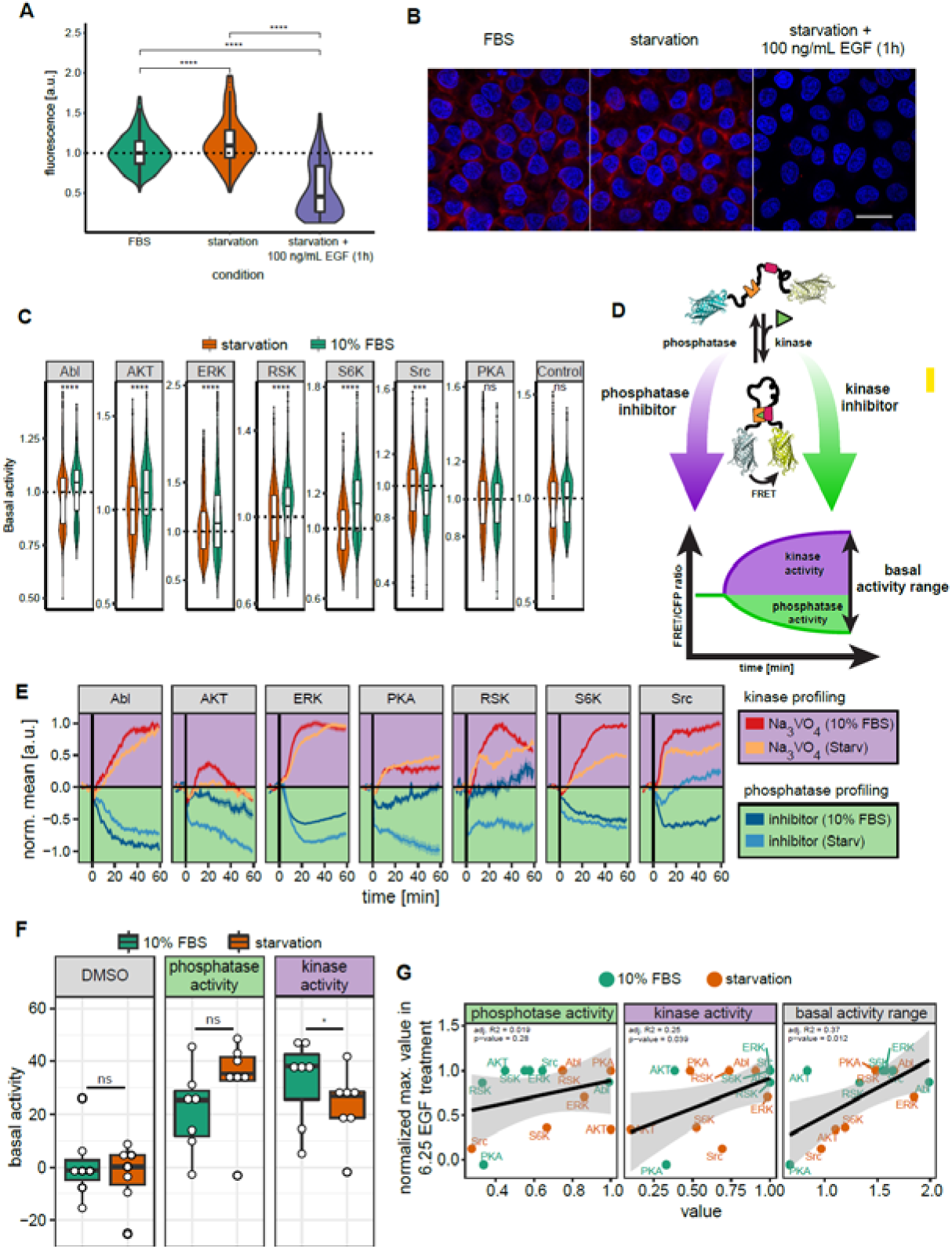
Extracellular environment fine-tunes basal kinase and phosphatase activity to tune signaling responsiveness. (A) Quantification of the EGFR surface abundance by staining with an antibody against the extracellular EGFR epitope. The fluorescent intensity (y axis) under various conditions (x axis) is depicted. Fluorescence intensities of the receptor were normalized to those in FBS. (B) Representative cell images from (A). Scale bar = 20 µm. (C) Basal activity profile in serum starved overnight and non-starved cells (kept in FBS constantly) acquired with FRET biosensors. P values are derived from Wilcoxon test. ns: p > 0.05, *: p ≤ 0.05, **: p ≤ 0.01, ***: p ≤ 0.001, ****: p ≤ 0.0001. n > 1000 individual cells per sensor and condition. (D) Illustration of the experimental setup to study kinase and phosphatase basal activity. (E) Quantification of kinase and phosphatase basal activity upon tyrosine phosphatase or kinase inhibition in serum starved and non-starved cells. Cells expressing were treated with 10 µM multi-AGC kinase inhibitor (AT13148; for AKT, PKA, RSK, S6K biosensors), 10 µM MEK inhibitor (AZD6240; for ERK biosensor), 10 µM Src/Abl dual inhibitor (SKI-606; for RSK and ABL biosensors) or 100 µM protein-tyrosine phosphatase inhibitor (activated Na_3_VO_4_; all biosensors). Mean ± SEM is shown. (F) Basal global kinase and phosphatase activity measured by FRET sensors in absence or presence of 10% FBS. The area under the curve were calculated after DMSO or inhibitor treatment in (E) as the approximation of enzyme activity. (G) Correlation of the phosphatase (left), kinase activities (middle) and basal activity range (right) with sensor responsiveness (normalized maximum value) after 6.25 ng/ml of EGF treatment. Shaded area represents 95% confidence intervals.

To investigate how the interplay between multiple extracellular cues might cooperate to mediat distinct signaling states, we took advantage of those FRET biosensors that are able to monitor the balance between kinase and phosphatase activity^55^. We also applied inhibitors to dissect the underlying mechanisms of tuning basal signaling activity by using FRET sensors as readout of kinase and phosphatase activity (**Figure 5D**). Upon treatment of serum starved cells with sodium orthovanadate (Na_3_VO_4_), a general protein-tyrosine phosphatase inhibitor^56^, we confirmed that the majority of biosensors including EGFR, ERK, RSK, S6K, SRC, and FAK exhibited elevated FRET ratios supporting the fact that kinase signaling is active in the absence of ligand, but is continually repressed by phosphatase activity in living cells^57^ (**Figure S5D**). Importantly, we did not observe AKT kinase activity upon starvation. In addition to the changes in kinase biosensors, we detected elevated activity of RhoA and Cdc42 GTPases in the presence of orthovanadate (**Figure S5D**). We identified that AKT, RSK, S6K, and SRC phosphorylated FRET biosensors more efficiently in the presence of 10% FBS compared to serum starved cells **Figure S5D**). In contrast, JNK and PKA were slightly less active in the presence of serum. Together, these results suggest that the basal activity of tyrosine kinases is under tight control of a complex extracellular environment.

We further profiled basal phosphatase activity that is regulating the phosphorylation level of the SRC, AKT, S6K, RSK, ERK, ABL, and PKA FRET biosensors by inhibiting kinase activities. The SRC/ABL dual inhibitor (SKI-606) induced a stronger dephosphorylation level of the SRC and ABL biosensors in the presence then in the absence 10% FBS, suggesting a higher activity of phosphatases compared to starved cells (**Figure 5E**). In contrast, treatment with the multi-AGC kinase inhibitor AT13148 as well as the MEK inhibitor AZD6240 resulted in a stronger dephosphorylation of the AKT, S6K, RSK, PKA, and ERK FRET biosensors upon serum starvation than in the presence of 10% FBS (**Figure 5E**). Overall, the kinase and phosphatase activity profiling identified that global kinase activity was increased in the presence of 10% FBS in comparison to starved cells indicating stronger basal activity of GF signaling network in the presence of FBS (**Figure 5F**).

The complex interplay of kinase and phosphatase activity can result in increased sensitivity of the two-component signal transduction system^58^. To account for the simultaneous effect of basal kinase and phosphatase activity, we calculated the basal activity range in serum starved and non-starved cells (**Figure 5D**). We detected a strong increase in basal activity by the ABL, S6K, and SRC FRET biosensors in the presence of 10% FBS. In contrast, the AKT and PKA FRET biosensors reported an increased basal activity range upon starvation (**Figure S5E**). Importantly, we have observed stronger correlation between the basal activity range and the signaling response to the low concentration of growth factors in comparison to phosphatase and kinase activities alone (**Figure 5G**). This observation suggests that the responsiveness of cells to the low concentration of growth factor depends on the interplay of kinase and phosphatase activities. Collectively, these results demonstrate that the extracellular environment regulates the basal phospho-turnover rate of the signaling network. More broadly, our results reveal that information from a signal-rich environment can be encoded by basal kinase and phosphatase activity to fine-tune a response and sensitivity to a newly incoming cue.

## Discussion

In this study, we used a FRET biosensor-based multiparameter imaging platform (FMIP) to investigate how cells integrate signals from multiple input cues. Our FMIP-based assay revealed a diverse landscape of signaling responses, including antagonism, additivity, and synergy, following stimulation of distinct receptor groups (**Figure 6** and **2E**). At a systems level, we demonstrated that these time-dependent signaling signatures were unique to each condition and were influenced by ligand concentrations. Notably, we uncovered a complex network of signaling crosstalk between components of distinct pathways, where the interplay of extracellular cues dictated signaling outcomes (**Figure 2F**). While we cannot exclude the potential role of receptor trafficking in modulating network activity, our data strongly support the notion that extracellular cue combinations and their concentrations fine-tune cellular signaling through intricate crosstalk among network components.

**Figure 6.**
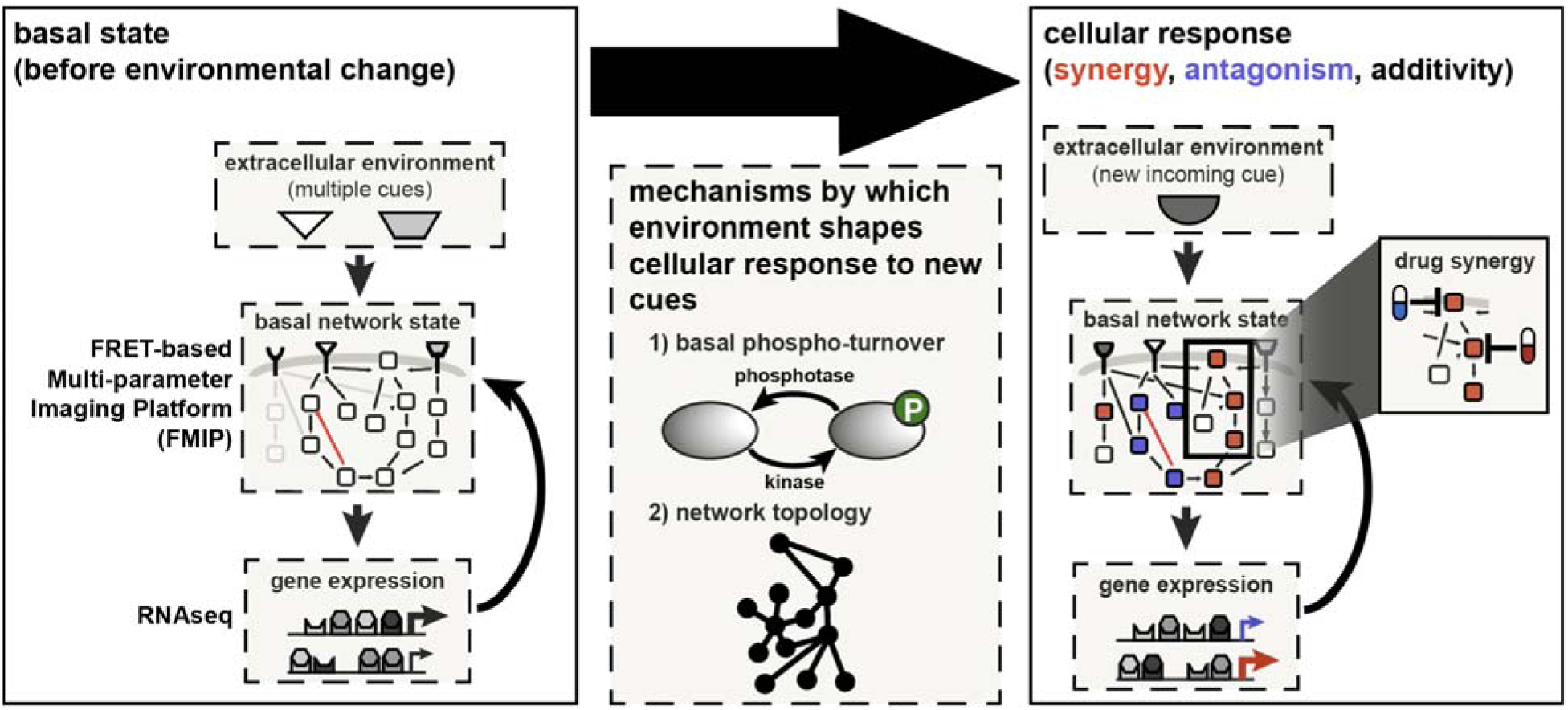
Activity-based network model of signal processing in response to environmental changes. Network activity is regulated by protein abundance, basal phosphorylation states, and network topology. The signaling network encodes information about the extracellular environment through dynamic phospho-turnover and topology-driven protein interactions. Changes in the extracellular environment can modulate these parameters, thereby tuning the network’s sensitivity and dynamic response to newly incoming signals. Consequently, signaling outputs are adjusted in an environment-dependent manner, exhibiting both linear and non-linear integration signatures—such as additivity, synergy, and antagonism. These dynamic signatures provide a functional readout that can be leveraged to identify synergistic drug combinations.

The identification of distinct signaling signatures raises an important question about the biological and clinical relevance of these interaction modes. The physiological relevance of this crosstalk was evident in perturbation experiments available in a public database. As predicted by signaling signatures with drug combinations, disrupting cross-talk synergistically impacted cellular survival and proliferation (**Figure 2I**). Moreover, signature-base node-node interactions were able to effectively identify synergistic drug combinations in comparison to signaling based node-node interactions (**Figure 2I**). Thus, profiling signaling signatures might be a more effective strategy to predict synergistic drug combinations than analyzing signaling dynamics alone.

Our RNA-seq analysis revealed that each signaling network state corresponded to a unique gene expression profile, as well as distinct transcriptional signatures of antagonism and synergy (**Figure 6, 3B and S3C-F**). Notably, we found that antagonistic and synergistic interactions at the RNA expression level were associated with distinct cellular processes, suggesting an additional layer of regulatory control over cellular physiology (**Figure 3F**). Our findings in HeLa cells align with previous studies across multiple cell types and species, where strong combinatorial interactions at the transcriptional level have been observed in response to growth factors and cytokines^15,17–19,59^. While we identified a large number of concentration-dependent interactions in gene expression following pairwise EGF and IGF treatments, many appeared weak at very low growth factor concentrations. Therefore, we anticipate that integrating more complex combinatorial treatments with physiologically relevant, low ligand concentrations will yield biologically meaningful effects on gene expression.

Cells are constantly exposed to multiple cues under physiological conditions. Surprisingly, under quasi-physiological conditions, mimicked by the presence of 10% of FBS, cells are more sensitive to lower concentration of growth factor. The balance of kinase and phosphatase activities plays a critical role in signal transduction^60^. Previously, Kleiman et al. identified that phosphatase activity persists under growth factor stimulation of high concentration. It was proposed that high rates of the activation/deactivation cycle may increase responsiveness to a drug treatment and allow kinetic proofreading^61^. Our experiments using kinase and phosphatase inhibitors while monitoring kinase and phosphatase activities over time suggested that the balance between protein tyrosine phosphatase and kinase activity may be of greater significance in cellular adaptation to the changing environment than so far realized. We suggest that phospho-turnover allow the basal network state to encode information on past experience of a cell and in turn adjust response at the signaling network level to the newly incoming signals through signaling signatures (**Figure 6**). In support of this hypothesis, we showed that under quasi-physiological conditions, in the presence of multiple signaling cues at low concentrations (namely in the presence of serum or GF combinations), the cellular signaling network is pre-activated and tuned to achieve specific sensitivity and dynamic response to a newly incoming cue (**Figure 4B-E and S4C**) of low concentration. Such sensitivity and dynamic response to a newly incoming cue are programed by the basal signaling state and encoded in network topology (**Figure 3F**) and phospho-turnover (**Figure 5E-G**). Basal activity range that combines kinase and phosphatase activity has higher prediction power of signaling response than any of the two alone (**Figure 5G**). It suggests that altered signaling program in response to a low concentration of EGF (**Figure 4B-E**) stems from distinct turnover rates of phosphorylation sites. Notably, we show that non-linear signaling signatures are very dynamic (**Figure 2E**) and have impact on gene expression (**Figure 3A**). Overall, our data supports that a network topology and phospho-turnover allow the signaling network machinery to integrate information from past experience and newly incoming cues that result in unique physiological response of the cell.

The current prevailing view is that aberrant signaling programs under pathological condition are predominantly induced by mutations or large changes in protein abundance^48,62–65^. However, other recent studies show that the extracellular microenvironment dramatically changes in diseased versus normal conditions^1,66–68^. Such changes in the microenvironment through autocrine and paracrine GF/cytokine signaling may result in altered signaling dynamics^69,70^. In this cellular context, the kinase-phosphatase balance could have far-reaching consequences for development of new treatment strategies and understanding how cancer cells adapt to the dynamic extracellular environment. For example, it is generally well established that the activation or amplification of the c-Met receptor tyrosine kinase is able to induce drug resistance in EGFR family mutant cancers^71–74^. However, the driving mechanisms leading to resistance are unclear. Intriguingly, we showed that low concentrations of HGF exhibited a strong synergistic effect on global EGF and IGF-1 signaling (**Figure 2G**). Thus, it is tempting to speculate that specific signaling signatures (such as synergy) might be a driving force in the selection of beneficial sub-clones in response to therapeutic pressure and have clear implications for drug development.

In conclusion, our methodology supports a comprehensive activity-based network model (Figure 6) of signal processing in the presence of multiple extracellular cues and provides a rich resource for future studies. Further studies based on this methodology will likely unveil novel mechanisms of cell adaptation to environmental changes including the evolution of drug resistance in the context of cancer.

## Author contribution

C.S. and D.K. designed the study, D.K. performed most of the experiments and the statistical analysis, F.Z., D.K., and W.H. performed statistical analysis of the RNA-seq data, F.S. performed immunoblot analysis. A.T. conducted experiments with growth factor pre-treatment. M.C. performed proliferation and cell cycle analysis. D.K. and C.S. wrote the manuscript with suggestions from all authors. U.K. contributed critically on the level of the study. C.S. and U.K. provided material support. The project was led by D.K. and supervised by C.S.

## Star Methods

### Key resources table

| REAGENT or RESOURCE | SOURCE | IDENTIFIER |
| --- | --- | --- |
| Antibodies |  |  |
| Mouse anti-EGFR | Santa Cruz Biotechnology | Sc-101, RRID: AB_627494 |
| Rabbit anti-EGFR | Cell Signaling | 4267, RRID: AB_2246311 |
| Rabbit anti-c-Abl | Cell Signaling | 2862; RRID: AB_2257757 |
| Rabbit anti-Src | Cell Signaling | 2108; RRID: AB_331137 |
| Rabbit anti-pEGFR (Tyr1068) | Cell Signaling | 2234, RRID: AB_331701 |
| Rabbit anti-pS6 (Ser235/236) | Cell Signaling | 2211, RRID: AB_331679 |
| Mouse anti-beta-Actin | Sigma-Aldrich | A5441, RRID: AB_476744 |
| Mouse anti-Vinculin | Santa Cruz | sc-55465, RRID: AB_630433 |
| Alexa Fluor 568 goat anti-mouse IgG (H+L) | Molecular Probes | A-11031, RRID: AB_144696 |
| Chemicals, Peptides, and Recombinant Proteins |  |  |
| Dulbecco's Modified Eagle's Medium | Sigma-Aldrich | 31885-023 |
| Fetal Bovine Serum | Sigma-Aldrich | F7524 |
| Fetal Bovine Serum | Gibco | 10270 |
| MEM | Sigma-Aldrich | M3024 |
| Lipofectamine 2000 Transfection Reagent | ThermoFisher Scientific | SKU#11668-019 |
| DRAQ5 | Cell signaling | 4084 |
| Src/Abl inhibitor SKI-606 | Selleck Chemicals | S1014 |
| EGF | Sigma-Aldrich | E9644 |
| TGF-α | Sigma-Aldrich | T7924 |
| TNF-α | Sigma-Aldrich | T6674 |
| IGF-1 | Sigma-Aldrich | I3769 |
| HGF | R&D systems | 294-HG-005 |
| Wnt-5a | R&D systems | 645-WN/CF |
| Gas6 | R&D systems | 885-GSB |
| PDGF-BB | Sigma-Aldrich | P4306 |
| H<sub>2</sub>O<sub>2</sub> | Sigma-Aldrich | H1009 |
| TRIzol reagent | Life Technologies | 15596-018 |
| SlowFade Diamond antifade mountant with DAPI | ThermoFisher Scientific | S36968 |
| Experimental Models: Cell Lines |  |  |
| H838 | U. Klingmueller | N/A |
| HeLa Kyoto | R. Pepperkok | N/A |
| Recombinant DNA |  |  |
| EKAREV FRET sensor | M. Matsuda (Komatsu et al., 2011) | N/A |
| ERK2G1 FRET sensor | O. Pertz (Fritz et al., 2013) | N/A |
| Ras Eevee FRET sensor | M. Matsuda (Komatsu et al., 2011) | N/A |
| Rac1 Eevee FRET sensor | M. Matsuda (Komatsu et al., 2011) | N/A |
| RSK Eevee FRET sensor | M. Matsuda (Komatsu et al., 2011) | N/A |
| S6K Eevee FRET sensor | M. Matsuda (Komatsu et al., 2011) | N/A |
| Akt Eevee FRET sensor | M. Matsuda (Komatsu et al., 2011) | N/A |
| PKC Eevee FRET sensor | M. Matsuda (Komatsu et al., 2011) | N/A |
| JNKAR1EV FRET sensor | M. Matsuda (Komatsu et al., 2011) | N/A |
| AKAR3EV FRET sensor | M. Matsuda (Komatsu et al., 2011) | N/A |
| PIP3/PI(3,4)P2 FRET sensor | M. Matsuda (Ananthanarayanan et al., 2005) | N/A |
| Src | Y. Wang (Ouyang et al., 2008) | N/A |
| Cdc42 Eevee FRET sensor | M. Matsuda (Kuchenov et al., | N/A |
|  | 2016) |  |
| PAK1 FRET sensors | C. Parrini (Parrini et al., 2009) | N/A |
| Calcineurin FRET Sensor | J. Zhang (Newman and Zhang, 2008) | N/A |
| FAK FRET Sensor | Y. Wang (Seong et al., 2011) | N/A |
| Prin-cRaf FRET sensor | M. Matsuda (Terai and Matsuda, 2005) | N/A |
| Prin-bRaf FRET sensor | M. Matsuda (Terai and Matsuda, 2005) | N/A |
| EGFR/abl FRET sensor | M. Matsuda (Komatsu et al., 2011) |  |
| Histone H3-K9 methylation FRET sensor | A. Ting (Lin et al., 2004) | N/A |
| Histone H3-K27 methylation FRET sensor | A. Ting (Lin et al., 2004) | N/A |
| Histone H3-S28 phosphorylation FRET sensor | A. Ting (Lin and Ting, 2004) | N/A |
| Cdk1 FRET sensor | J. Pines(Gavet and Pines, 2010) | N/A |
| PDK1 FRET sensor | J. Zhang (Gao et al., 2011) | N/A |
| Caspase-3 FRET sensor | P. Sorger (Albeck et al., 2008) | N/A |
| Caspase-8/10 FRET sensor | P. Sorger (Albeck et al., 2008) | N/A |
| PKC FRET sensor | A. Newton (Violin et al., 2003) | N/A |
| Rap1 FRET sensor | M. Matsuda(Mochizuki et al., 2001) | N/A |
| RalA FRET sensor | M. Matsuda (Takaya et al., 2004) | N/A |
| RhoA FRET sensor | M. Matsuda (Yoshizaki et al., 2003) | N/A |
| TN-XL (Ca<sup>2+</sup>) FRET Sensor | O. Griesbeck (Mank et al., 2006) | N/A |
| Twitch (Ca<sup>2+</sup>) FRET Sensor | O. Griesbeck (Thestrup et al., 2014) | N/A |
| Pickles (Abl) FRET sensor | Y. Ohba (Mizutani et al., 2010) | N/A |
| Aurora B FRET sensor | M. Lampson (Fuller et al., 2008) | N/A |
| Control FRET | S.Vogel (Koushik et al., 2006) | N/A |
| Control (DEVG) FRET | P. Sorger (Albeck et al., 2008) | N/A |
| Pyruvate FRET sensor | L. Barros (San Martin et al., 2014) | N/A |
| Glucose FRET sensor | W. Frommer (Takanaga et al., 2008) | N/A |
| DAG FRET sensor | C. Schultz (Kuchenov et al., 2016) | N/A |
| Software and Algorithms |  |  |
| R |  | https://www.r-project.org |
| Fiji | (Schindelin et al., 2012) | https://fiji.sc |
| FluoQ | (Stein et al., 2013) | N/A |
| DESeq2 | (Love et al., 2014) | N/A |
| Pscan | (Zambelli et al., 2009) | http://159.149.160.88/pscan/ |
| Other |  |  |
| Drying pearls Orange | Sigma-Aldrich | 94098 |
| Nunc Lab-Tek | ThermoFisher | 155361 |

## Supplemental information

**Figure S1.**
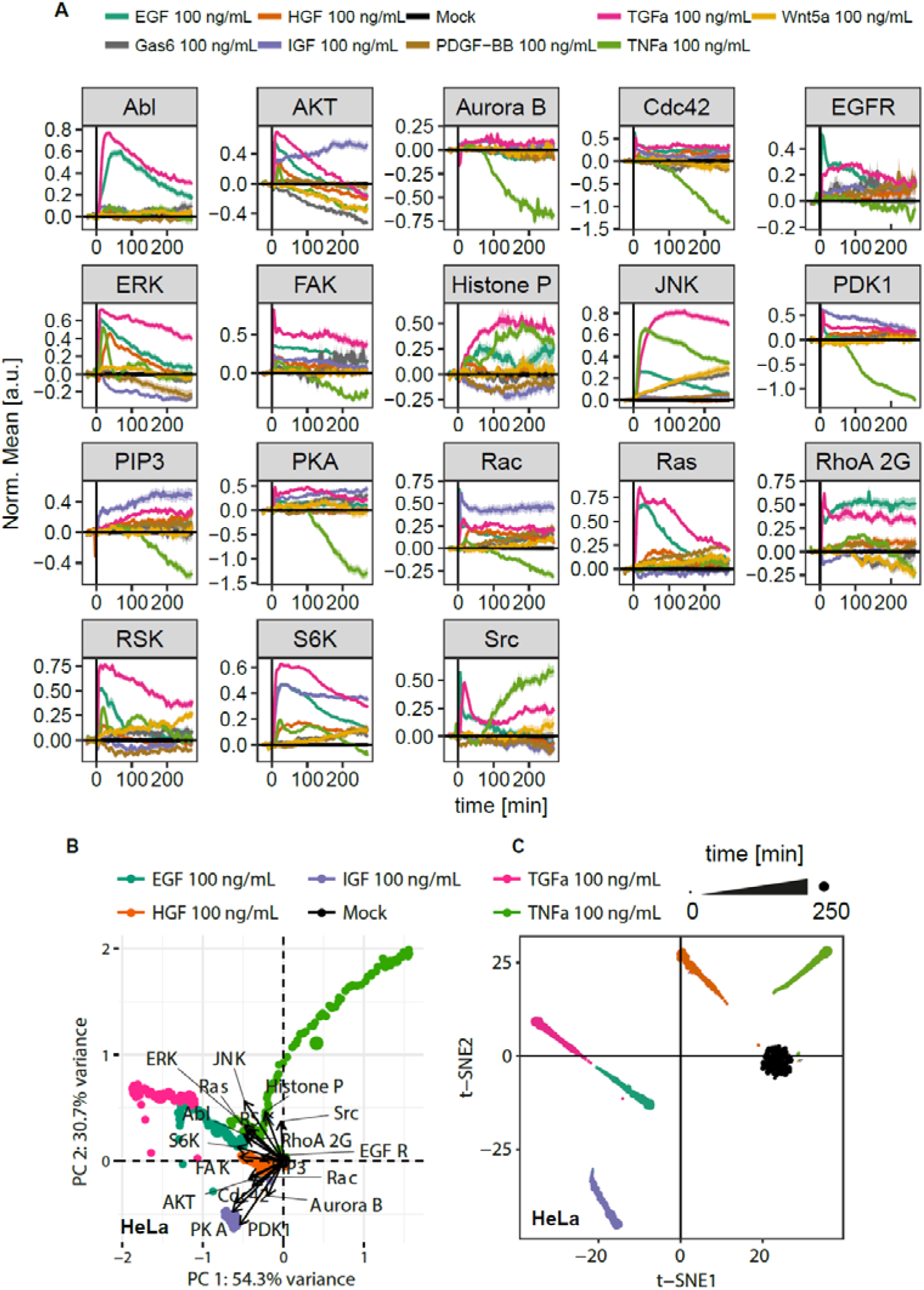
related to Figure 1. Growth factor/cytokine signaling in HeLa cells. (A) FRET biosensors responded to growth factor and cytokine stimulation. HeLa cells were stimulated with 100 ng/mL EGF, 100 ng/mL IGF-1, 100 ng/mL HGF, 100 ng/mL TGFα, 100 ng/mL TNFα, 100 ng/mL PDGF-BB, 100 ng/mL WNTa or 100 ng/mL Gas6 at time 0. Data represent mean *±* SEM (n ≥ 2). (B) Principal component analysis (PCA) of the HeLa signaling state after EGF, TGFα, IGF-1, HGF, TNFα and control treatment. The PCA scores (points) and loadings (vectors) correspond to the samples and FRET sensors, respectively. 18 loading vectors indicating the impact of individual FRET sensors. Each dot represents a single time point. (C) t-SNE analysis of the average response to growth factors/cytokines. Each dot represents a single time point.

**Figure S2.**
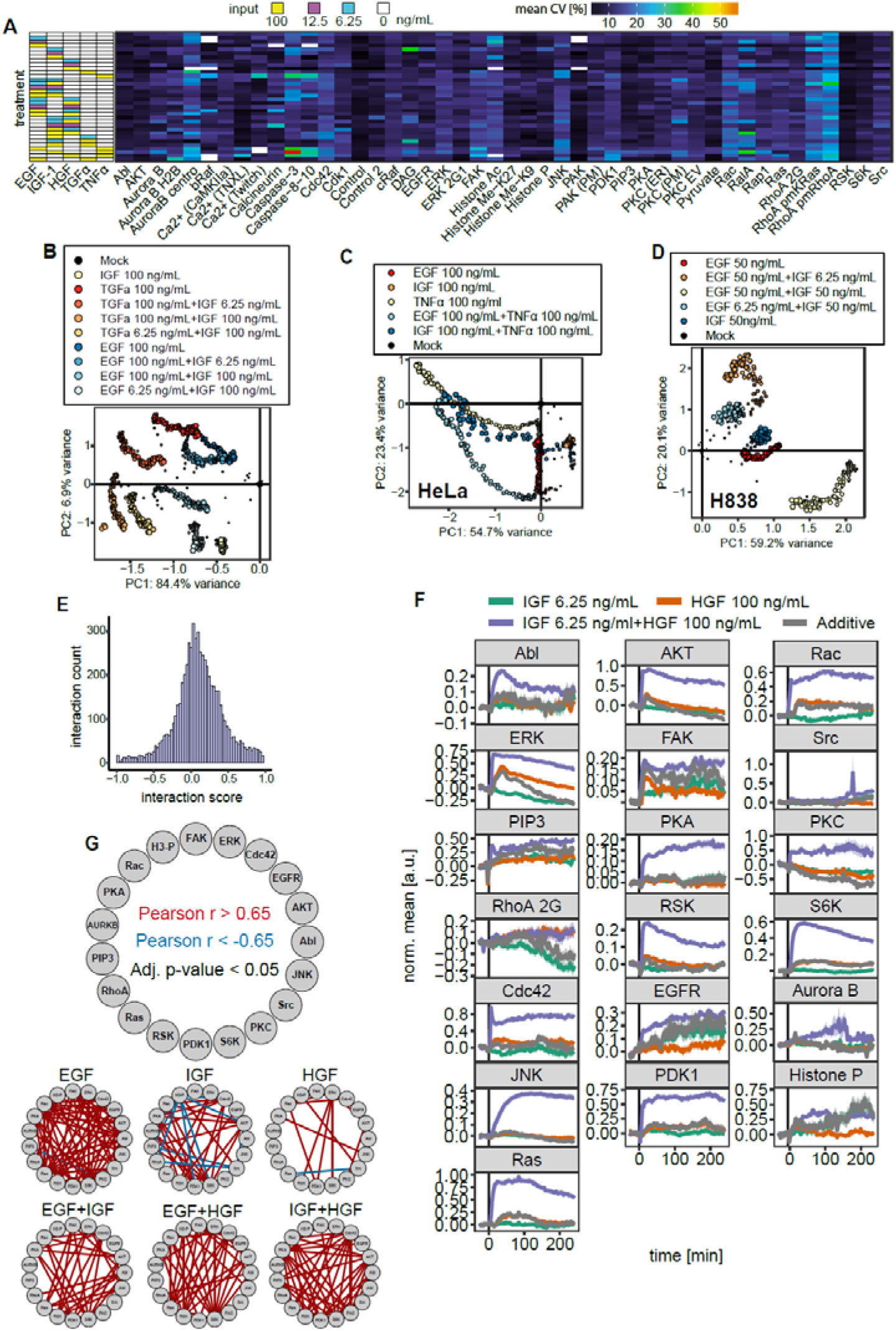
related to Figure 2. GF signaling interactions in living cells. (A) Heatmap showing the CV across single-cell responses was calculated at each time point under each stimulation (along the side) for each of the FRET biosensors (along the bottom). Th mean of the resultant CV values is shown. (B) Comparison of global signaling response for TGFα/IGF-1 and EGF/IGF-1 using PCA. Each dot represents a single time point. The size of the dot illustrate time after treatment in min. (C) Global signaling response to the combinations of two ligand pairs projected into the first two principle components. HeLa cells were stimulated with 100 ng/mL EGF, 100 ng/mL IGF-1, 100 ng/mL TNFα, 100 ng/mL EGF + 100 ng/mL TNFα, 100 ng/mL IGF-1 + 100 ng/mL TNFα or untreated. (D) Global signaling response to the combinations of three ligand pairs projected into the first two principle components. H838 cells were stimulated with EGF, IGF-1 or their combinations. (E) Distribution of interaction scores for pairs of growth factors across all FRET biosensors and time windows. (F) Difference between calculated (or expected) and experimental response. Mean ± SEM are shown. (G) Graph representations of strong Pearson correlations between signaling events for EGF, IGF-1, HGF, EGF/IGF-1, EGF/HGF and HGF/IGF-1. Red and blue edges represent positive (r > 0.65 with adjusted p-value < 0.05, Student’s t test) and negative (r < -0.65 with adjusted p-value < 0.05, Student’s t test) correlations, respectively.

**Figure S3.**
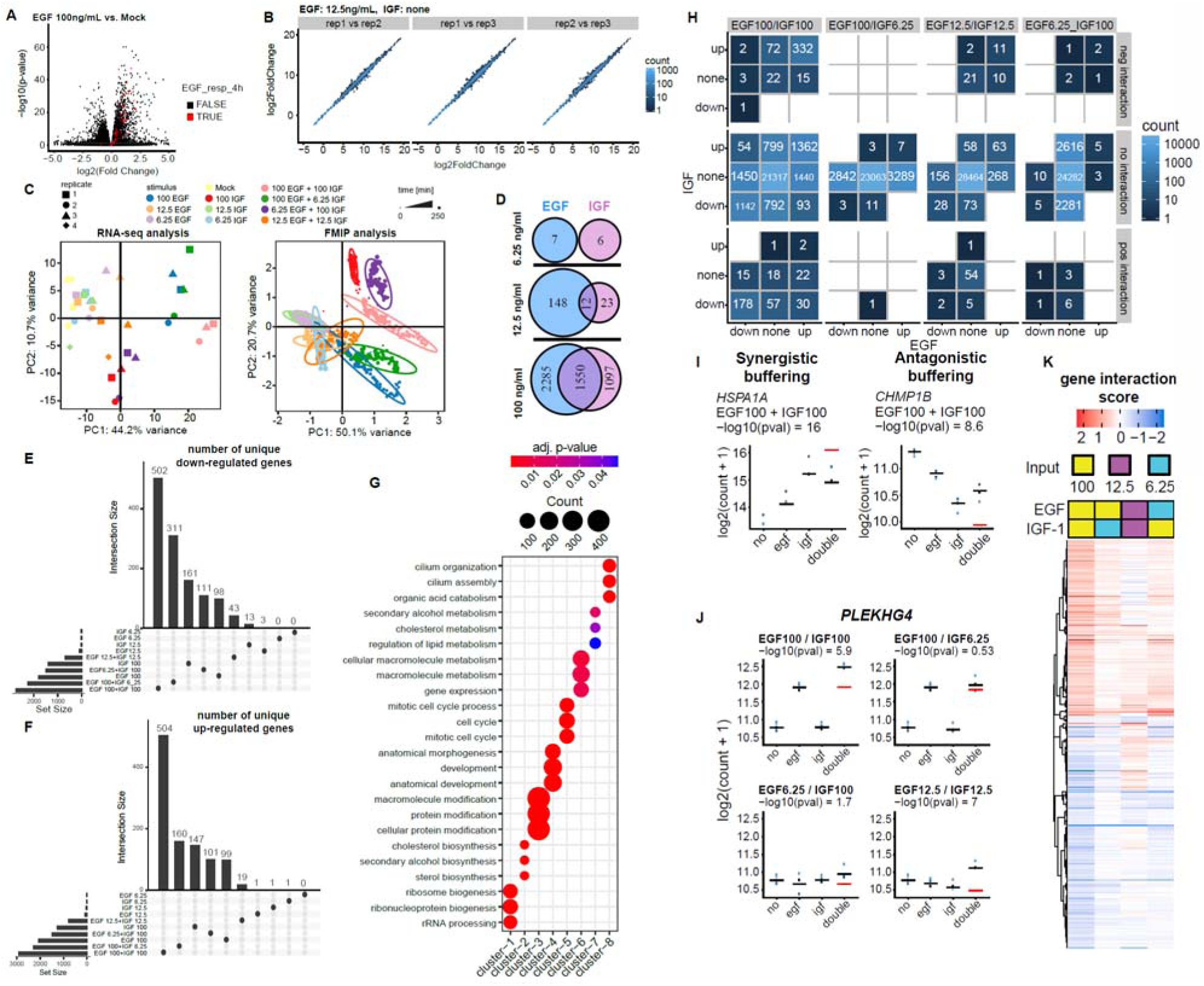
related to Figure 3. Pairwise growth factor treatments profoundly affect global gene expression profile. (A) Comparison of genes regulated by an EGF treatment in a previously published study (Amit et al., 2007) to the 100ng/ml EGF treatment in our study. Red: Genes whose expression peaked in (Amit et al., 2007) at 240 min after stimulation of HeLa cells with 20ng/ml EGF. (B) Examples of global gene expression correlation for independent biological replicates in HeLa. (C) PCA analysis of RNA-seq data (left) and FMIP data (right). PCA of RNA-seq data is based on 10000 genes (independent biological replicates ≥ 3, n=4 for controls and n=3 for stimulated cells). PCA of FMIP data is based on FRET biosensor data measured by FMIP over 4 hours. Color indicates GF treatment. (D) Venn diagram portraying EGF or IGF-1 genes and their different response under the various concentrations (adjusted *p*-value ≤ 0.05, measured with DESeq2). (**E-F**) UpSetR (Conway et al., 2017) plot highlights the unique number of genes down-regulated (E) and up-regulated (F) across all conditions after 4h treatment with GF combination or individual growth factor. (G) Gene Ontology Biological Process (GOBP) enrichment analysis of genes in each of the clusters identified in (Figure 3C). The three most statistically significant GOBP terms for each cluster are depicted. Significant terms are marked as dots. The number of genes is indicated by the size of the dot. (H) Table showing the classification of the interaction response genes (IRGs) induced by the EGF/IGF-1. The figure classifies for the four different EGF/IGF combinations (four columns) whether a gene was significantly up, down or neither up nor down (“none”) regulated in the respective single treatment and subsequently classified whether the interaction for the combined treatment was significantly positive, negative or neither significantly positive nor negative. For example, in the EGF100/IGF100 treatment combination, there were 332 genes with a negative interaction where additional the EGF100 treatment and the IGF100 treatment show an up-regulation. (I) Expression of example genes classified in synergistic or antagonistic buffering. Red bar indicates expected expression upon pairwise treatment. (F) Expression of example genes across under various concentrations and ratios of EGF/IGF-1. Red bar indicates expected expression upon pairwise treatment. (**K**) Heatmap representation for clustering of gene interaction scores for all synergistic or antagonistic genes that responded to at least one pairwise treatment (FDR = 0.1).

**Figure S4.**
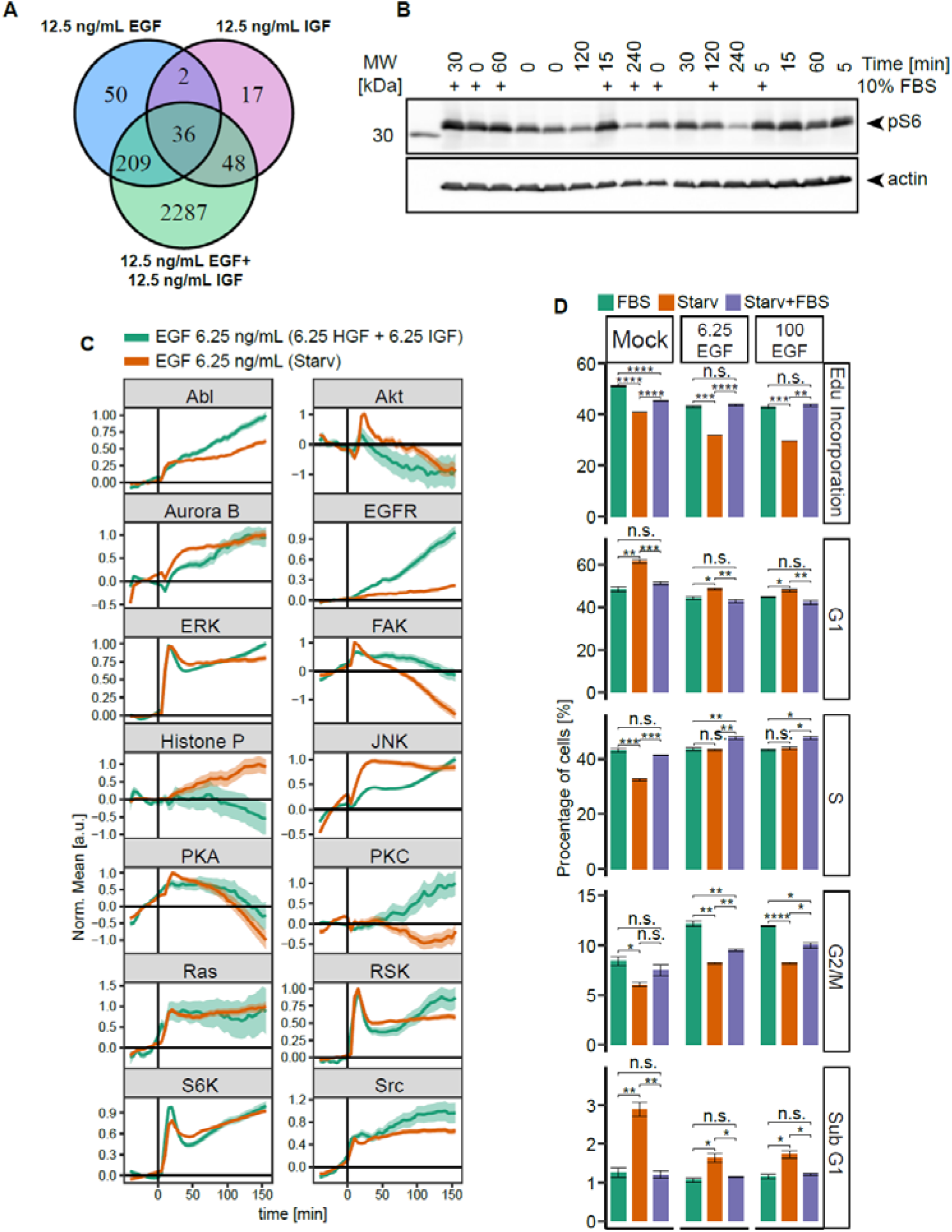
related to Figure 4. GF-induced genes, signaling and phenotypic changes. (A) Venn diagram portraying cytokine-induced different response under various treatments (adjusted p-value ≤0.05, measured with DESeq2). (B) HeLa cells were either growth factor depleted or maintained in 10 % FBS and stimulated with 6.25 ng/ml EGF. Phosphorylation of S6 was detected by quantitative immunoblot. An exemplary immunoblot is shown. Sample loading was randomized to avoid systematic errors. (C) Quantification of signaling activity after exposure to 6.25 ng/mL EGF in absence or presence of 6.25 ng/ml IGF-1+ 6.25 ng/ml HGF. HeLa cells were starved or incubated with 6.25 ng/ml IGF-1+ 6.25 ng/ml HGF overnight. Curves indicate mean ± SEM (n = 3). (D) Comparison of 5-bromodeoxyuridine (EdU) incorporation and each cell cycle (G1, S and G2/M) phase measured by 4,6-diamidino-2-phenylindole (DAPI) staining for cells kept in full media (green), serum starved cells (orange) or starved cells treated with 10% FBS (violet). Cells were unstimulated or stimulated with 6.25 ng\ml or 100 ng\ml EGF. Error bars represent ± SEM. Significance was assessed by Student’s t test. ns: p > 0.05, *: p ≤ 0.05, **: p ≤ 0.01, ***: p ≤ 0.001, ****: p ≤ 0.0001.

**Figure S5.**
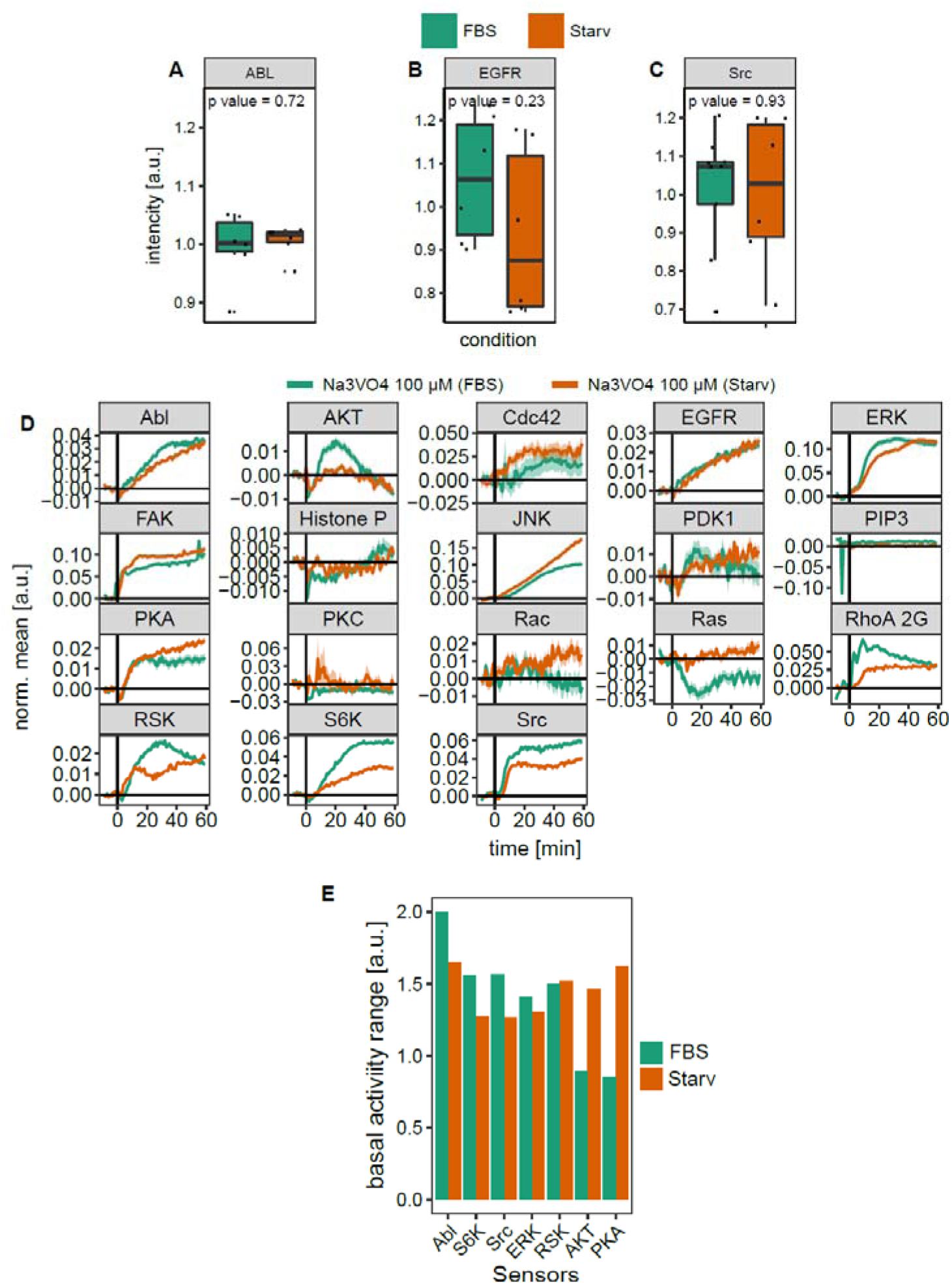
related to Figure 5. FBS does not change expression of ABL, Src and EGFR and modulates signaling activity and basal activity range. (**A-C**) Box plot representation of ABL (A), EGFR (B) and Src (C) protein levels for cells kept in full media or serum starved cells measured by western blot analysis. (D) Distinct signaling activity upon tyrosine phosphatase inhibition in serum starved and non-starved cells. Cells were treated with 100 µM Na_3_VO_4_. Mean ± SEM are shown. (E) Quantification of basal activity range from (Figure 5E)

## Resource availability

### Lead contact

Further information and requests for resources and reagents should be directed to and will be fulfilled by the Lead Contact Carsten Schultz.

### Cells and reagents

H838 cells were passaged in high glucose DMEM (Life Technologies) supplemented with 10 % fetal bovine serum (FBS, Sigma) and with 100 units/mL penicillin, and 100g/mL streptomycin (Pen/Strep, Invitrogen). HeLa Kyoto cells were maintained in low glucose DMEM (Life Technologies) supplemented with 10 % FBS and with 100 µg/mL of primocin (Invivogen). Starvation of H838 cells was performed in high glucose DMEM (Lonza) supplemented with 1 mg/mL of BSA (Sigma), 2mM L-glutamine (Invitrogen), 100 units/mL penicillin, and 100g/mL streptomycin. Starvation media for HeLa Kyoto cells contained low glucose DMEM (Life Technologies) and 100 µg/mL of primocin (Invivogen). The dual Src/Abl inhibitor (SKI-606, **Bosutinib**), the MEK inhibitor (AZD6244) and the multi-AGC kinase inhibitor (AT13148) were purchased from Selleck Chemicals. DMSO was obtained from Merck KGaA. EGF, IGF-1, TGF-α, PDGF-BB and TNF-α were obtained from Sigma. HGF, Wnt-5a and Gas6 were purchased from R&D systems.

### Contact printing

The fabrication of microarrays was performed as described previously in (Kuchenov et al., 2016). Plasmids for reverse transfection were diluted to concentration of 1 mg/mL. The transfection mixture was prepared by mixing 9 µl of a 0.4 M sucrose solution in DMEM, 9 µl of DNA and 33µL of lipofectamine 2000 mixed in a 96-well plate and incubated for 20 min at room temperature. Subsequently, 21.75 µL of 0.29% gelatin solution in water was added to the transfection mixture. The transfection cocktail was distributed in 384-well plates (24 µL per well). The plates were stored at -20°C. In order to print Labteks the 384-well plate was thawed at room temperature and centrifuged briefly up to 54 g to straighten the surface of the samples. Afterwards, residual bubbles were removed by a 10 µL tip and placed immediately in the contact printer. Before printing, LabTek dishes were washed with 70% ethanol increasing hydrophobicity of the LabTek surface and, accordingly, improving the shape of the spots. 1-well LabTek dishes were printed with a “ChipWriter” contact printer equipped with solid pins. Using PTS 600 pins, the diameter of printed spots was about 600 µm and the spot-to-spot distance was 1125 µm. Printed 1-well LabTek dishes were stored at room temperature in a gel drying box in the presence of drying pearls (Sigma).

### Live cell imaging

HeLa (650,000) or H838 (800,000) cells were transiently transfected by seeding on printed LabTeks. The seeding was performed when cells were cultured to 50%-60% confluence. After maintaining HeLa and H838 cells in an incubator for 24 or 48 hr, respectively, the media were changed to starvation media 12–17 hr prior to imaging. During imaging, cells were maintained in imaging medium (minimum essential medium supplemented with 100 U/mL of penicillin, 100 mg/mL of streptomycin, and 30 mM of HEPES) at 37 C without CO2. To assist cellular segmentation cells were preincubated with 7.5 nM DRAQ5 for 30 min (Cell Signaling Technology).

Time-lapse imaging was performed on an Olympus IX83 microscope equipped with a Hamamatsu ImagEM CCD camera and an environmental control unit incubation chamber using 20x 0.70 numerical aperture (NA) or 10x 0.40 NA. The microscope was equipped with 436/20 excitation filter, a CFP/yellow fluorescent protein (YFP) dualband beam splitter (51017bs; Chroma), and two emission filters (470/30 for CFP and 535/50 for YFP) that were controlled by a filter wheel. Image acquisition was controlled by the Olympus IX83 microscope software – xCELLence (Olympus) or cellSens (Olympus). Fluorescence images were acquired with an exposure time of 200 ms.

### RNA extraction and sequencing

HeLa cells were seeded in 60 mm dishes pre-coated with gelatin to reproduce conditions similar to microarrays. Prior to stimulation cells were starved in the absence of serum overnight. Afterwards, cells were incubated with GFs for four hours. We lysed cells by the direct addition of 3 ml of TRIzol (Life Technologies) after media removal according to the manufacturer’s instructions. For RNA extraction lysates were treated with 0.6 ml chloroform (Sigma-Aldrich) for 3 min and centrifuged at 12,500 g for 15 min at 4 °C. Aqueous supernatant was collected and diluted 1:1 with 70% ethanol. Total RNA was extracted from solution using RNeasy Mini Kit (Qiagen), following the manufacturer’s instructions and quantified using the NanoDrop spectrophotometer. RNA was used with A(260/280) nm ≥ 1.8 and A(260/230) nm ≥ 2.0. RNA quality was assessed using RNA 6000 Nano chips on the Agilent 2100 Bioanalyzer. The library preparation, RNA sequencing and reads alignment were performed by a Genomics Core Facility at EMBL. Sequencing was performed on Illumina NextSeq-500 instruments.

### Immunofluorescence

HeLa cells were seeded in a 12-well glass-bottom dish (Ibidi, 81201) and maintained in CO2 incubator for 24h. Afterwards, the medium was changed to full or FBS-free medium. After, 17h of incubation, cells were fixed with 4% paraformaldehyde (PFA; Electron Microscopy Sciences, 19208) in PBS for 10 min at room temperature and washed with PBS. Cells were blocked in 1% (w/v in PBS) BSA for 1 h at room temperature. After removal of the blocking solution (without any wash), cells were incubated with the anti-EGFR primary antibody (Santa Cruz Biotechnology, sc-101, 1:200) diluted in 1% (w/v, in PBS) BSA at 4 °C overnight. After washing with 1% (w/v, in PBS) BSA, cells were incubated with goat anti-mouse Alexa Fluor 546 (Life Technologies, A-11031) at room temperature for 1 hour. Cells were then washed with PBS at room temperature and finally rinsed with distilled water. After drying, coverslips were covered with SlowFade Diamond antifade mountant with DAPI (ThermoFisher Scientific, S36968). Microscopy images were captured at room temperature using a confocal laser scanning microscope (Zeiss LSM780) with a 63× oil objective and fully opened pin hole. Settings were as follows: DAPI-channel: 405-nm excitation (ex), 409- to 475-nm emission (em); red channel: 561-nm ex, 569- to 655-nm em. Images were further processed using Fiji software (fiji.sc/) with the in-house developed macro.

### Quantitative immunoblot analysis

For the analysis of phosphorylated and total S6, total EGFR and total Abl, 6·10^5^ HeLa cells were seeded one day in advance in a 4 cm cell culture dish (TPP) and washed three times with DMEM without additives and then incubated for 16 hours in growth factor depletion medium or growth medium containing 10 % FBS. The cells were stimulated with the indicated doses of EGF (GF144, Millpore) for the given timespan. The cells were lysed and processed as previously described (Merkle et al., 2016). Cell lysates were separated by SDS-PAGE and transferred to a PVDF membrane (0.45 μm pore size, Immobilon, Millipore). Antibodies against phosphorylated S6 (#2211), total EGFR (#4267), total Src (#2108) and total Abl (#2862) were obtained from Cell Signaling. Antibodies for loading control were purchased from Santa Cruz (anti-vinculin, sc-55465) and Sigma (anti-actin, A5441). Secondary antibodies coupled to hrp were obtained by Dianova. Luminescence was detected by ImageQuant LAS4000 (GE Healthcare) and signal intensity was quantified by ImageQuant TL software (Version 7.0, GE Healthcare).

### EdU incorporation assay

HeLa Kyoto cells were seeded into two 15 cm dishes at 20% confluency in DMEM supplemented with 10% FBS (DMEM + 10% FBS). 24 hours later, one dish (A) was washed once with DMEM only, and then serum-starved in DMEM only for 16 hours. The other dish (B) was washed once with DMEM + 10% FBS, and then cultured in DMEM + 10% FBS for 16 hours. At the end of the 16-hour incubation, cells on both dishes were detached using TrypLE Select (1X) and collected by centrifugation. Cells from A were washed once with DMEM only, and then re-suspended in DMEM only at 2 x 10^5^ cells/mL density. Cells from B were washed once in DMEM supplemented with 10% FBS, and then re-suspended in DMEM + 10% FBS at 2 x 10^5^ cells/mL density. 1 mL of suspensions from A and B were added to 35 mm dishes that already contained 1 mL of 2X concentration of the growth factors prepared in DMEM only and DMEM + 10% FBS, respectively. 24 hours later, 20 µL of EdU (1 mM) in PBS was added into each dish (final concentration 10 µM) for a 2-hour period. One dish was not treated with EdU and used as negative control. At the end of EdU labelling, cells were washed with PBS, detached from the dishes using TrypLE Select (1X), washed by centrifugation with PBS, and then fixed in 70% ethanol at +4 °C overnight. The next day the ethanol was removed, and then a copper-catalyzed click reaction was performed using Alexa 488 - picolyl azide supplied in Click-iT® Plus EdU Alexa Fluor® 488 Flow Cytometry Assay Kit from Thermo Fisher (#C10632). Cells were counterstained using DAPI (10 µg/mL final conc.), and finally analyzed for proliferation and cell cycle by BD LSRFortessa (BD Biosciences) cell analyzer.

### Image analysis

The single image of a nuclei channel and the time-lapse series of CFP and FRET channels acquired from different positions were exported as separate TIFF files. The images were analyzed with FIJI (Schindelin et al., 2012) and FluoQ (Stein et al., 2013). The FIJI-based macro developed in-house was used to preprocess and multiply nuclei image and subsequently concatenate all channels together. This macro produced a single tiff file containing three channels: binary mask of nuclei (repeated 100 times), FRET and CFP channels obtained from the same position. The file containing three channels was further analyzed with the FIJI-based macro FluoQ. Although cells express different amount of FRET biosensors we used image analysis pipeline that automatically account for a low expressing cells that are close to background cellular fluorescence. In order to subtract background we used a histogram-based “Triangle” algorithm to calculate the mean of the thresholded background that was subsequently subtracted from each pixel. Next, image smoothing with a median filter (radius size = 2) was applied and the images were transformed to a 32-bit float. Cells were automatically segmented by using binary mask created by Huang’s fuzzy thresholding method. The signal intensity that was equal or close to intensity of the background was set as NaN value due to Huang’s fuzzy thresholding. This image analysis pipeline automatically accounts for a very low expressing cells and avoided erroneous FRET ratios. The nuclear binary mask was used to segment cell by the voronoi algorithm. In order to define ROIs the particle analyzer, a build-in FIJI plugin, was applied to the binary image of segmented cells. Although acquired images contain information of the subcellular activity we simplified image analysis pipeline in order be able to analyze 13900 images (more than 5000 cells) from a single experiment in a reasonable time window by averaging the intensity of FRET, CFP and FRET ratio over each ROI. However, significant compartmentalized signaling fluctuations might be averaged out. In all experiments, single-cell traces were normalized to the of the FRET ratio from before stimulation. The output file in a text format that was produced by FluoQ contained all measured parameters, statistical summaries.

### Outliers removal

In each independent experiment, single-cell traces were cautiously examined for artificial intensity spikes in the CFP and FRET channels. Those spikes were due to lamp intensity fluctuation, cell division, cell movement and, in rare cases, cell death. Therefore, we have developed the automated algorithm that detected artificial intensity spikes and removed cell traces containing those spikes from further data analysis. First, the baseline and response values were smoothed by running median (width of median window = 5). Subsequently, the difference between real values and smoothed values was used to calculated sample quantiles. The quantile mean and standard deviation for both baseline and response was calculated from the values belonging to the interval lying between 5^th^ and 95^th^ quantiles. Afterward, the z-score was calculated for each value of the baseline and response using quantile mean and standard deviation. Single cell traces containing z-score in the experimentally defined range (z-score ≥ 40 or z-score ≤ -40) were automatically removed. After removal single cell traces containing spikes, the mean baseline (formula 1), the standard deviation of the baseline (formula 2), the slope of the baseline (formula 3), the mean (formula 4), maximum (formula 5) and minimum (formula 6) response were calculated for each single-cell trace. Those features were subjected to the

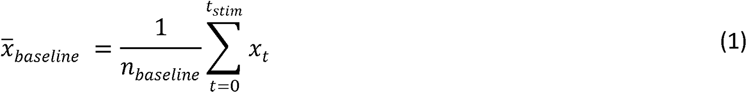

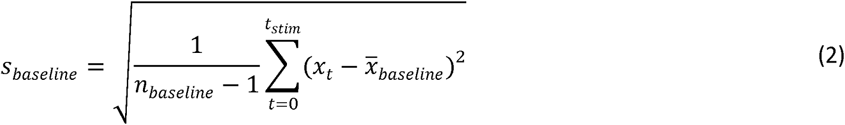

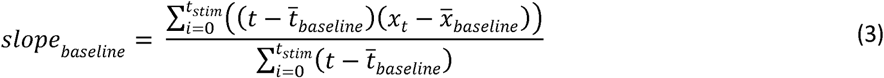

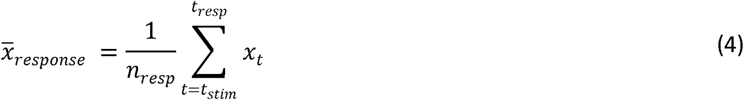

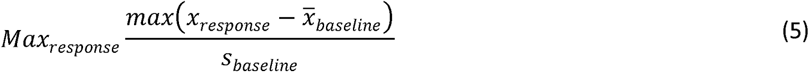

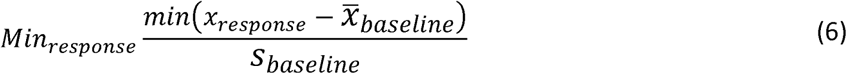

algorithm that classifies an outlier if it falls outside the interval defined in the following formula:

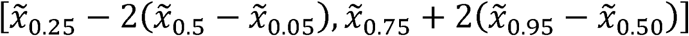

Once any feature of a particular single-cell trace was detected as an outlier, the cell was excluded from the analysis.

### Interaction Score

In order to calculate an interaction mode between two stimuli for each signaling node (or FRET biosensor), single-cell traces first were normalized by dividing each time point with the mean baseline (*x̅ baseline*) and with the mean value of untreated cells (x_mock) at each time point followed by a subtraction of 1. The resulted single-cell trace was normalized to the maximum observed average value across all conditions (formula 7). Afterwards, area under the curve (AUC) was computed for each single-cell trace within a specified time window (formula 8).

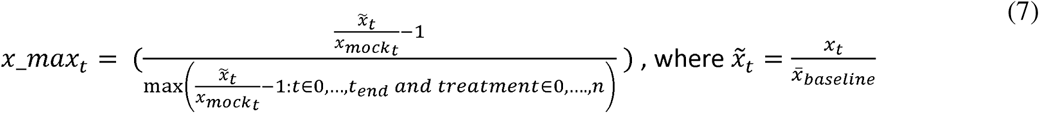

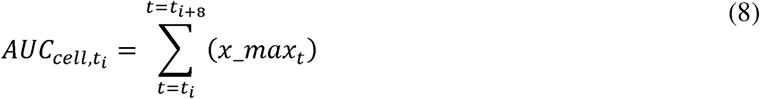

The expected response upon co-stimulation was calculated by the simple addition of the AUC mean of individual treatments. The combined error was estimated by an error propagation of the individual standard errors of the mean applying a Variance formula (Ku, 1966). In order compare the experimental and expected (calculated) additivity the means of two groups were subjected to Student’s two sample t-test and subsequently obtained P-values were corrected for multiple testing (Benjamini and Hochberg, 1995). We defined the ‘interaction score’ as the difference between the experimental and expected (calculated) additivity. In order to simplify visualization, the interaction score was scaled to the maximum interaction score observed for a FRET biosensor, giving a value that ranges from −1 (antagonism) to +1 (synergy). Adjusted p value < 0.05 was considered to be statistically significant.

### Analysis of RNA-seq data

RNA-seq reads were aligned with salmon version 0.8.2 (Patro, 2017) to the Human Reference Genome GRCh38 obtained from Ensembl (http://www.ensembl.org). Differential expression analysis was performed using the R/Bioconductor package DESeq2 (Love et al., 2014). To account for systematic effects between biological replicates, a blocking factor was added to the design formula prior to differential testing. Genes with an FDR < 0.05 were considered significant.

Interaction analysis was carried out by supplying DESeq2 with a custom design matrix. The columns of the matrix correspond to factor levels of the respective EGF and IGF concentration and replicate as well as four columns corresponding to the interaction of the combined EGF/IGF treatment for the concentrations EGF:100ng/ml+IGF:100ng/ml, EGF:100ng/ml + IGF:6.25ng/ml, EGF:12.5ng/ml + IGF:12.5ng/ml and EGF:6.25ng/ml+IGF:100ng/ml respectively. Interaction scores were computed as model coefficient corresponding to the respective interaction column in the design matrix. Enrichment analysis was performed for genes with significant interactions using Fisher’s exact test with the R package piano (Varemo et al., 2013) on the Hallmark (Liberzon et al., 2015) gene set collection of MSigDB (Subramanian et al., 2005). Throughout the analysis, the false discovery rate (FDR) for calling differential expression or interaction was controlled using the method of Benjamini and Hochberg (Benjamini and Hochberg, 1995). If not otherwise stated, gene interaction scores with an FDR < 0.1 were considered significant.

### Transcription factor binding sites analysis

Lists of 119 synergistic and 88 antagonistic genes (|GIS| >0.5 and adjusted p-value <0.05) were uploaded to Pscan website (http://159.149.160.88/pscan/). The JASPAR non-redundant database (Jaspar 2018_NR) (Khan et al., 2018) and the promoter region between −450 and +50 bp of gene were used for motif analysis.

### General statistical analyses and FRET data visualization

Single-cell traces first were normalized by dividing each time point with respect to the mean baseline (*x̅ baseline*) and with respect to the mean value of untreated cells (x_mock) at each time point followed by a subtraction of 1. The resulted single-cell traces were normalized to the maximum observed average value across all conditions (formula 7). The treated starved cells were normalized to starved “Mock” cells whereas treated cells kept in 10% FBS were normalized to untreated “Mock” cells maintained in 10% FBS. Unless otherwise stated, a FRET biosensor response is represented as normalized FRET ratio mean of all individual cells from identical conditions +/- SEM where the SEM is calculated from all cells in all experiments with identical conditions.

### Principal component analysis (PCA) and t-SNE analysis

The prcomp function without centering and scaling of the program R was used to perform PCA. The Rtsne package was used to perform t-SNE analysis. PCA and t-SNE was performed on X x Y matrix, where Y is different FRET biosensors each with X that included: time points, stimuli and stimuli doses.

### Computation of correlation networks based on FMIP dataset

To compute correlation networks, we first combined interaction score values across all time points, concentrations and ratios in a single vector for each FRET biosensor. For each growth factor pair, we used the Pearson correlation coefficient to quantify correlation between each pair of FRET biosensors. Correlation analyses were performed in R with Benjamini-Hochberg multiple testing correction separately within each condition. The correlation matrices were calculated by using the ‘rcorr’ function of Hmisc package (Harrell, 2016) and visualized by using R corrplot package. Network diagrams were generated using the igraph package (Csardi and Nepusz, 2006).

### Compilation of drug-drug interaction dataset and statistical analysis

We extracted the drug-drug interaction scores from DrugcombDB database (version: May 2019) (ref). HSA and ZIP scores were selected for further analysis because their averaged score across whole database is close to 0 value, indicating that initial dataset is not biased towards synergistic/antagonistic drug-drug combinations. Additionally, on this data, we integrated the data that associates each drug with experimentally identified or computationally predicted target(score ≥ 850) extracted from DrugcombDB database. Resulting dataset contained 84349 unique drug combinations and 123 cell lines. We further selected drug combinations that specifically target nodes classified in four categories: 1) whole database; 2) drug combinations targeting node-node interactions identified by both signaling- and signature-based approaches; 3) drug combinations targeting unique signaling-based node-node interactions; 4) drug combinations targeting unique node-node interactions identified by signature-based approach. We used the two-tailed unpaired t test to determine if there was a significant difference between selected categories.

### Comparison of FRET sensor and RNAseq datasets using co-inertia analysis

The co-inertia analysis (CIA) (Doledec and Chessel, 1994) was conducted utilizing the ADE-4 package, a versatile tool designed for multivariate statistical analysis widely employed in environmental and ecological data analysis. This approach aimed to elucidate the primary associations between the RNAseq expression and FMIP signaling profiles within each specific condition, using HeLa cells as the cell line model. Given the time series nature of the FRET-based dataset, a timepoint aggregation within 4 hour window was performed for each condition by summing all individual timepoints together. CIA was performed on two X x Y matrices, where Y is different conditions (stimuli and stimuli doses) for both FMIP and RNAseq datasets. X in FMIP dataset represents different FRET sensors, whereas X in RNAseq dataset represents different genes.

### Correlation analysis between FMIP and RNAseq datasets

Classical Pearson correlation, which represents the linear relation between two variables was used in this study to identify relationship between responses of FRET sensor response and gene expreesion. Pearson correlation was computed using the stats R package. Given the time series nature of the FRET-based dataset, a timepoint aggregation within 4 hour window was performed for each condition by summing all individual timepoints together (an approximation for area under the curve). The correlation matrices were calculated by using the ‘rcorr’ function of Hmisc package (Harrell, 2016)

### Clustering of correlation data between FMIP and RNAseq datasets

The genes showing significant correlation between FRET sensor response and gene expression (adjusted p value < 0.05) for one or more sensors were selected. Selected genes and clustered using hierarchical clustering (complete-linkage) with euclidean as the distance measure.

## Data visualization

If not stated otherwise, the data was visualized using the R package ‘ggplot2’ (Wickham, 2009). The overall FRET ratio for a reporter is represented as a normalized FRET ratio mean of all individual cells from identical conditions ± SEM, where the SEM is calculated from all cells in all experiments under identical conditions.

